# De novo mutations across 1,465 diverse genomes reveal novel mutational insights and reductions in the Amish founder population

**DOI:** 10.1101/553214

**Authors:** Michael D. Kessler, Douglas P. Loesch, James A. Perry, Nancy L. Heard-Costa, Brian E. Cade, Heming Wang, Michelle Daya, John Ziniti, Soma Datta, Juan C Celedón, Manuel E. Soto-Quiros, Lydiana Avila, Scott T. Weiss, Kathleen Barnes, Susan S. Redline, Ramachandran S. Vasan, Andrew D. Johnson, Rasika A. Mathias, Ryan Hernandez, James G. Wilson, Deborah A. Nickerson, Goncalo Abecasis, Sharon R. Browning, Sebastian Zoellner, Jeffrey R. O’Connell, Braxton D. Mitchell, Trans-Omics for Precision Medicine (TOPMed), TOPMed Population Genetics Working Group, Timothy D. O’Connor

## Abstract

*de novo* Mutations (DNMs), or mutations that appear in an individual despite not being seen in their parents, are an important source of genetic variation whose impact is relevant to studies of human evolution, genetics, and disease. Utilizing high-coverage whole genome sequencing data as part of the Trans-Omics for Precision Medicine (TOPMed) program, we directly estimate and analyze DNM counts, rates, and spectra from 1,465 trios across an array of diverse human populations. Using the resulting call set of 86,865 single nucleotide DNMs, we find a significant positive correlation between local recombination rate and local DNM rate, which together can explain up to 35.5% of the genome-wide variation in population level rare genetic variation from 41K unrelated TOPMed samples. While genome-wide heterozygosity does correlate weakly with DNM count, we do not find significant differences in DNM rate between individuals of European, African, and Latino ancestry, nor across ancestrally distinct segments within admixed individuals. However, interestingly, we do find significantly fewer DNMs in Amish individuals compared with other Europeans, even after accounting for parental age and sequencing center. Specifically, we find significant reductions in the number of T→C mutations in the Amish, which seems to underpin their overall reduction in DNMs. Finally, we calculate near-zero estimates of narrow sense heritability (h^2^), which suggest that variation in DNM rate is significantly shaped by non-additive genetic effects and/or the environment, and that a less mutagenic environment may be responsible for the reduced DNM rate in the Amish.

**Significance:** Here we provide one of the largest and most diverse human *de novo* mutation (DNM) call sets to date, and use it to quantify the genome-wide relationship between local mutation rate and population-level rare genetic variation. While we demonstrate that the human single nucleotide mutation rate is similar across numerous human ancestries and populations, we also discover a reduced mutation rate in the Amish founder population, which shows that mutation rates can shift rapidly. Finally, we find that variation in mutation rates is not heritable, which suggests that the environment may influence mutation rates more significantly than previously realized.

## Introduction

*de novo* mutations **(DNM)** appear constitutively in an individual despite not being seen in their parents, and their identification and study are critically important to our understanding of human genomic evolution^1-11^. For example, it is necessary to understand the rate at which DNMs accumulate in order to calibrate evolutionary models of species divergence. DNMs are also implicated in many diseases, including rare genetic disorders^8,12,13^, and common complex diseases, such as autism and schizophrenia^13-15^. Early studies indirectly inferred mutation rate estimates from patterns of rare Mendelian diseases or from DNA substitution rates between species^4,16,17^. More recently, modern sequencing technologies have enabled the use of pedigree data to directly estimate the number of new mutations found across the genome^17-19^. These pedigree based studies have identified both paternal and maternal age effects, and estimate contributions of 1.51 and 0.37 DNMs per year of paternal and maternal age, respectively^8,19,20^. While these effects are often explained on the basis of DNA replication errors, recent studies have found that DNA repair processes are likely to be major contributors to the mutations that accrue in both paternal and maternal gametes^21-24^. Some of this repair-associated mutation accumulation has been found to be due to maternal age dependent DNA damage in oocytes and to maternal age dependent post-zygotic mutation increases^23^. These studies have also identified specific mutation patterns, such as C→G transversions, that strongly associate with DNA double strand breaks and repair^23,24^. In addition, these repair-associated mutations have been found to cluster together in distinct patterns, to predominate in certain genomic regions, and to be associated with recombination^21,22,24^, which has itself been shown to influence mutation rates^25,26^. Other features have also been reported to influence variation in mutation rates across the genome. GC content was recently shown to directly increase single nucleotide and structural mutation rates, and was also interestingly shown to increase recombination rates^27^. Chromatin structure has also been shown to associate with DNA mutations, with DNA replication times associating significantly with point mutations^28^, and nucleotide positioning found to significantly modulate mutation rate^29^. Recent work has tied a number of these features together by providing resolute and individualized recombination maps, and using them to demonstrate a positive relationship between maternal age and the rates and locations of meiotic crossovers^30^. Specifically, older mothers have increased recombination that also shifts towards late replicating regions and low GC regions, and this study also identified numerous loci that seem to genetically influence meiotic recombination. Other recent work evaluates nucleotide content, histone and chromatin features, replication timing, and recombination to provide one of the clearest pictures to date of the factors that shape genome-wide variability in the human mutation rate^22^. With regard to base content, Amos^31^ has proposed the “Heterozygote Instability” hypothesis, which challenges the assumption that population size and mutation rate are independent, and suggests their interdependence on the basis of heterozygosity. According to this hypothesis, the occurrence of gene conversion events at heterozygous sites during meiosis could locally increase mutation rates, and Amos uses substitution rates to provide support for this^32^.

While identifying these mutational correlates have helped us better understand the biological processes that drive mutation, genetic estimates of mutation rate are one half the magnitude of those originally inferred phylogenetically^17^. This has raised questions about the accuracy of these genetics estimates, as well as about the accuracy of human evolutionary time points calculated using phylogenetic estimates. While it has been proposed that the failure of genetic methods to account for post-zygotic mutations in the parent might bias estimates down and partly explain this discrepancy^19^, recent work using sibling recurrence suggests only minor mutation rate estimate increases when accounting for a substantial portion of these mutations^33^. This discrepancy has also raised the possibility that mutation rates have evolved more rapidly than previously assumed, and that molecular clock type analyses are therefore flawed^17^. For example, analyses of base pair substitution patterns have identified mutational differences between human populations, and showed most notably that the rate of TCC→TTC transitions appears to have increased in Europeans thousands of years ago for some finite period of time^34,35^. This work demonstrated that mutational processes can change rapidly and have changed recently, and also suggested the possibility of identifying mutational modifiers.

These findings, along with Amos’ “Heterozygote Instability” hypothesis ^32^, provide a rationale for how mutation rates might differ between human populations. However, since most studies of DNMs have used data from small cohorts of individuals with predominantly European ancestry^8,10,36^, little is known about the role of DNMs in the evolution and health of populations of predominantly non-European ancestry, and it is unclear whether DNM rates vary across different human populations. To address this and other questions about mutation, we use a high-coverage whole genome sequencing dataset generated by the NHLBI Trans-Omics for Precision Medicine (TOPMed) program^37^ to directly estimate and analyze DNM accumulation across multiple human ancestries and populations. After analyzing genome-wide patterns of mutation using a call set of 86,565 single nucleotide variant (SNV) DNMs, we compare the number of SNV DNMs per individual across five TOPMed cohorts that represent European, African, and Native American (Latino) ancestry individuals, and that include Amish individuals from a founder population with European ancestry. We also estimate the correlation between heterozygosity and SNV DNM count, and then test whether mutation rate is a heritable trait in anticipation of using GWAS to look for mutation rate modifying loci. Finally, we explore whether environmental exposures and/or random effects might significantly shape mutation rates and processes.

## Results

### TOPMed dataset and positive correlation between DNMs and parental age

Using whole genome sequencing data for 1,465 individuals and their parents from the TOPMed initiative^37^, we identify a *de novo* mutation (DNM) call set and compare DNM accumulation rates across ancestral background (Table 1). The analyzed individuals belong to five TOPMed cohorts with varied ancestral backgrounds: the Amish are an an isolated European founder population, the Barbados Asthma Genetics Study (BAGS) consists of individuals with predominantly African ancestry, the Cleveland Family Study (CFS) consists of both European and African American individuals, the Genetic Epidemiology of Asthma in Costa Rica (GACRS) and the Childhood Asthma Management Program (CAMP) are collectively referred to as the CRA cohort and consist of admixed Latino individuals, and the Framingham Heart Study (FHS) consists of individuals with European American ancestry. For our analyses, we treated the CFS study as two separate cohorts (signified as CFS_EUR and CFS_AFR, respectively). These make up our six analysis cohorts (Amish, BAGS, CFS_EUR, CFS_AFR, CRA, FHS, Table 1). After removing samples with DNM counts that were extreme outliers (often due to pedigree errors, see methods), we were left with a DNM call set of 86,865 SNVs across 1,453 individuals. This equates to an average of about 60 mutations per individual, which is directly in line with previous findings^8,20,24^. These 1,453 individuals come from 1205 independent nuclear families, and for our analysis, we treat each child and there two parents as a trio. To control for any potential confounding effects from this sibling data, we repeat all of our analyses that focus on per individual DNM measures after randomly choosing one child per family. The results from these repeated analyses are qualitatively the same as those from our full analysis, and for simplicity, we only report results from analyses run with our full dataset (except when noted with our heritability models).

**Table 1.**
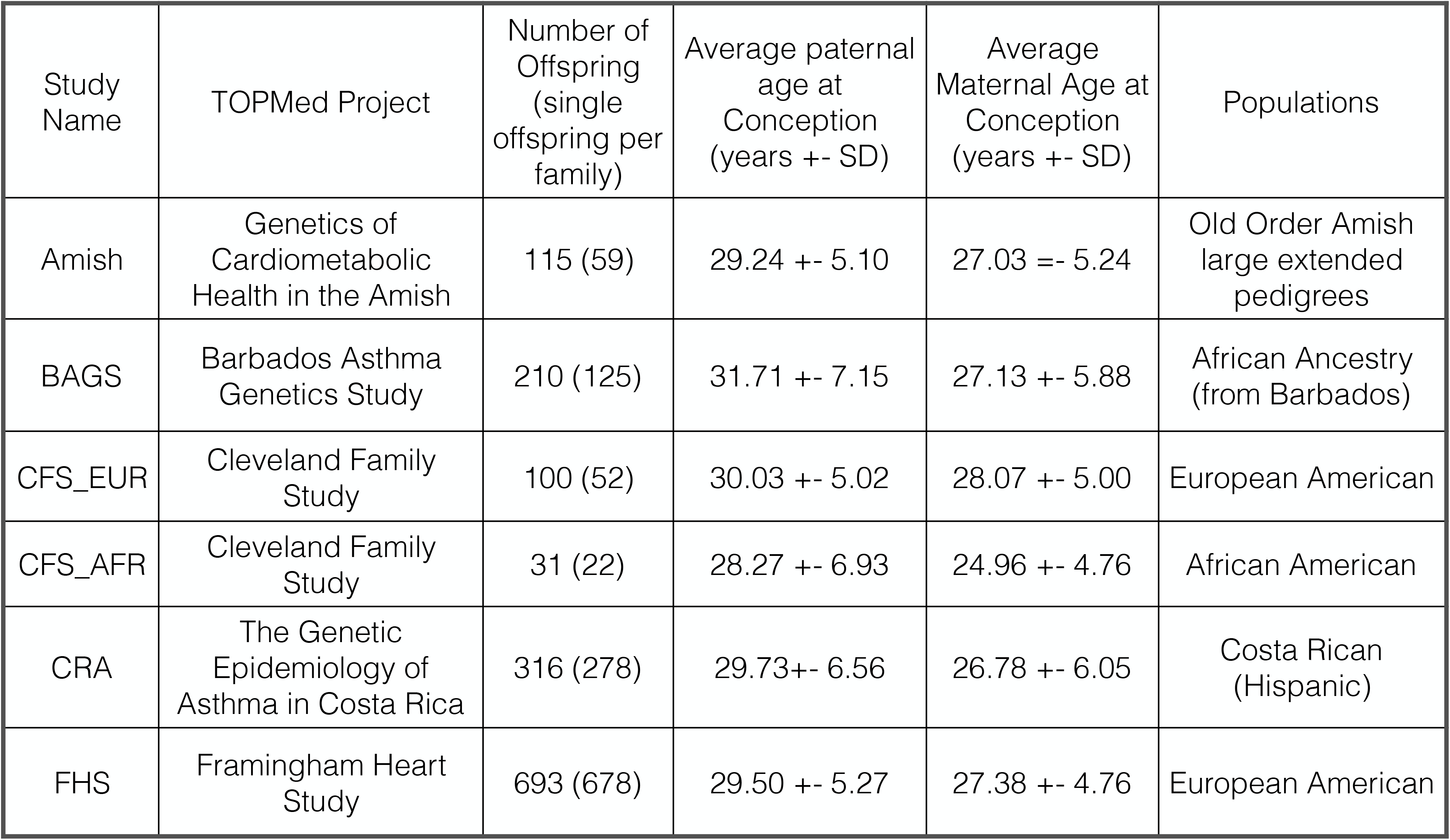
Study Characteristics. Study cohort and meta data are described. The six cohorts used in this study derive their names (“Study Name”) from 5 TOPMed projects (“TOPMed Project”), and represent a diversity of populations and ancestries (“Populations”). Sample sizes are shown (“Number of Offspring”), and average paternal and maternal ages are similar, with BAGS having the highest average paternal age, and CFS_AFR having the lowest maternal ages.

Employing this call set along with metrics from our sequencing experiments measuring coverage and quality, we estimate a human SNV DNM rate of 1.017 × 10^-8^ mutations per base pair per generation (see methods). This is in complete concordance with recent genetic estimates^17,38-40^, and lends support to our filtering approach. As expected based on previous studies^8,19^, we find a highly significant association between SNV DNM count per individual and paternal age (linear regression, R^2^=0.428, P<9×10^-172^, Figure S1A), which provides an additional degree of validation for our approach and our call set. While the high correlation in our dataset between paternal and maternal age makes it difficult to evaluate the separate effect of each on DNM count, a generalized linear Poisson model succeeds in identifying significant paternal and maternal age effects that are consistent in magnitude with those of recent studies (1.30 mutations per year of father’s age, and 0.37 mutations per year of mother’s age) ^19,20^. DNM totals per individual do not differ significantly on the basis of the sex of the individual for whom the DNMs are called (Figure S1B), their year of birth (Figure S1C), or the age of their DNA (i.e. the individual’s age) at the time of collection (Figure S1D).

### DNM mutation types and patterns

Consistent with previous findings, the most frequent mutation types in our DNM call set are C→T (43.47%) and T→C (25.53%) transitions (Figure 1A, Table S1). After accounting for the significant positive association between each DNM type and paternal age at conception (Figure 1A, blue stars), we identify significant positive correlations between maternal age at conception and C→T, C→G, and T→A mutations (Figure 1A, red stars). While DNM type composition is similar across the genome (Figure S2), regions with significant deviations from the mean may serve as good candidates for the identification and investigation of atypical mutational processes. When viewing each DNM as a 3-mer by considering the bases in the human reference genome immediately preceding and following each mutation, the influence of base context on mutational frequency can be better appreciated (Figure 1B). For example, T→C mutations preceded by an A appear to be more common than might have been predicted based on their central base pair mutation type alone, and they also comprise three of the twelve 3-mers significantly associated with paternal or maternal age at conception. While CpG to TpG transitions already comprise four of the five most common 3-mer DNMs, their mutational potential is particularly highlighted by normalizing each 3-mer DNM count for background 3-mer frequency, which demonstrates an excess of CpG to TpG transitions compared to the expectation based on genome frequency (Figure S3). Normalization also reveals other mutations with frequencies that are greater or lesser than what would have been expected on the basis of their central base pair mutation type (e.g. greater GCG→GAG, lesser TCT→TTT, etc), which further supports the notion that local base context influences mutational patterns.

**Figure 1.**
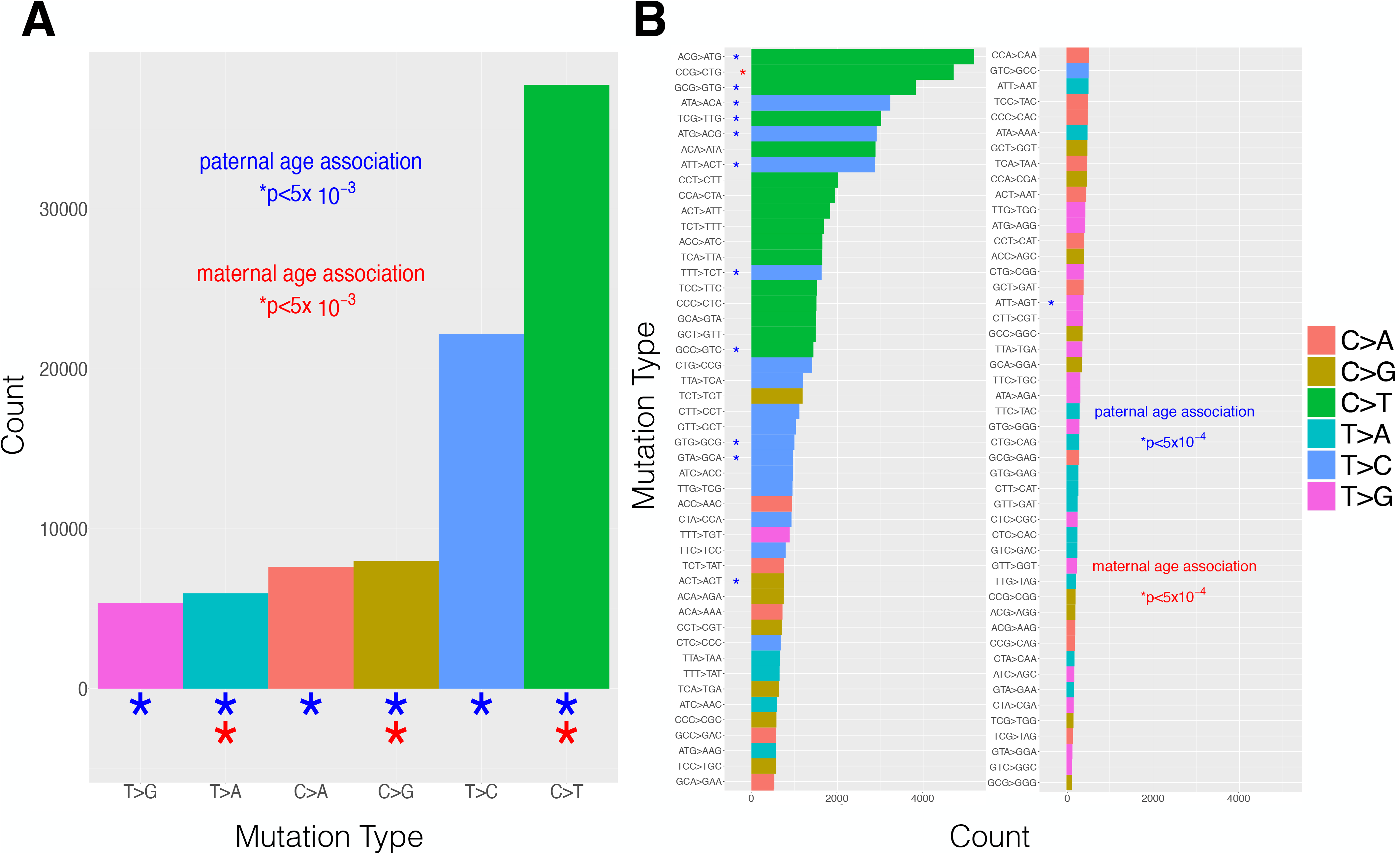
Distribution of single base and 3-mer mutation types across SNV DNM call set. A)The distribution of single base mutation type counts across our SNV DNM call set is shown. Colors represent mutation type, and stars represent associations with paternal (blue) and maternal (red) age (P < 5×10^-3^). B) The count across our SNV DNM call set for each of 96 3-mer mutation types is shown. Colors represent the center base mutation, and are the same as those in A. Stars represent associations with paternal (blue) and maternal (Red) age (P < 5.0 x10^-4^). P values for A and B were chosen to be more conservative than Bonferroni corrected values using an n of 6 and 96, respectively.

To evaluate the relationship between mutation and recombination, we use our DNM call set to calculate local mutation rate across 1 megabase (Mb) windows, and test the correlation between these rates and local recombination rate estimates from the International HapMap Consortium^41^. We find a significant positive correlation in which regions of higher recombination have more DNMs (Figure S4A, R^2^=0.0291, P=3.0 × 10^-19^). This relationship varies across the autosomes, with chromosomes 7, 8, and 16 showing strong positive correlations in which recombination explains as much as 35% of the variation in DNM rates (Figure S4B). Conversely, this correlation is either absent or limited across a number of the other autosomes. We then test the degree to which these local mutation rate estimates explain the distribution of genomic variation seen at the human population level. To do this, we use genomic variation data from the TOPMed consortium that was ascertained by performing whole genome sequencing on ∼41k diverse unrelated individuals^37^. We then calculate the total number of rare variants (AF < 0.005) within 1 Mb windows across the genome, and use normalized Z-scores derived from these counts as a measure of localized levels of recent human genomic variation. Notably, after accounting for local recombination rates, local DNM rates explain up to 30.79% of the variation in these normalized rare variation counts (P<1.5×10^-216^). When considering local recombination rates along with DNM rates, we can explain up to 35.48% of this rare genomic variation (P<9.5×10^-256^, Figure 2). Regions with the highest levels of rare variation, such as megabases 1-7 on chromosome 8 and 1-9 and 78-90 on chromosome 16, have the highest DNM rates, and are largely comprised of regions in the top 10% of recombination values. Interestingly, these regions have a unique DNM profile, with increased proportions of C→A and C→G mutations, and even greater proportions of C→T mutations than high recombination regions, which already have elevated C→T proportions compared to the genome average (Table S2). Conversely, low recombination regions have reduced C→T and C→G proportions, as well as increased proportions of T→A, T→C, and T→G mutations (Table S2, Figure S5). These patterns hold when using linear regression to implement a conservative adjustment for coding proportion and mappability concerns, with DNM rates still explaining 21.32% of the variation in rare variant levels (P<1×10^-141^), and DNM rates plus recombination rates explaining 27.3% (P<3×10^-186^, Figure S6). These associations reflect the degree to which *de novo* mutations shape the landscape of human genomic variation, and also have implications for the complex relationship between recombination and mutation.

**Figure 2.**
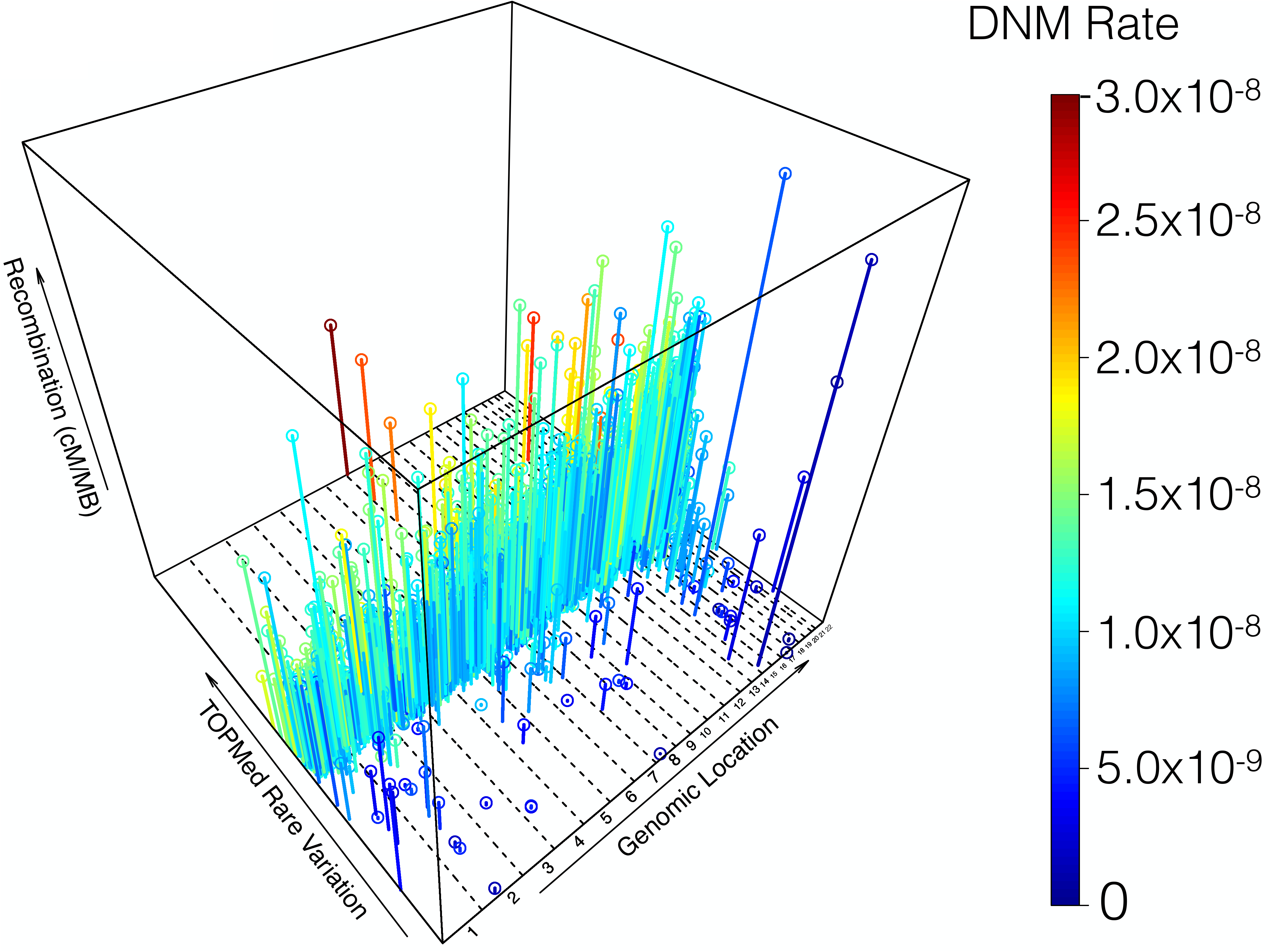
City Plot of rare variation, recombination rate, and DNM rate across the genome. The relationship between DNM rate (blue to red color range), rare variation (Y-axis, ranging from −5.83 to 9.06 Z-scores), and recombination rate (Z-axis, ranging from 0 to 7.77 cM/ Mb Y cM/Mb) across the genome (X-axis, dotted vertical lines divide autosomes 1-22). In going from low to high rare variation levels, a blue to red gradient can be seen, which reflects the significant correlation between DNM rate and population-level rare variation. Furthermore, regions with high DNM rates and high variation levels generally have taller bars, which reflects the positive relationship between DNM rate, variation level, and recombination (a few exceptions to this can be seen at the bottom corner of the plot as tall blue bars). Regions with the highest variation levels in the genome, such as those on chromosomes 8 and 16, have the highest DNM rates.

In testing the relationship between heterozygosity and DNM count across all individuals, we find a marginally significant positive correlation (P<0.01, Figure S7A). However, while this persists after adjusting for parental age (Figure S7B), heterozygosity at most explains 0.5% of the variation in DNM count. Furthermore, we find no relationship between heterozygosity and DNM count when looking at only individuals from FHS, which represents our largest cohort (Figure S8). This is only minimally consistent with predictions that heterozygosity drives *de novo* mutation, as the most recent work into the Heterozygote Instability hypothesis suggests a strong relationship between heterozygosity and DNMs^32^.

### DNM rate comparisons across ancestrally diverse cohorts

When comparing the number of DNMs per individual across all six ancestrally diverse cohorts (Table 1), we find significant differences, even after accounting for parental age (ANOVA, P<1.14×10^-3^, Figure 3,S9A). However, this difference appears to be driven by the Amish, who have significantly fewer DNMs per individual than the other cohorts. To control for sequencing center differences that could potentially influence the number DNMs identified, we performed a sub-analysis comparing DNMs between cohorts that were sequenced at the same center. This sub-analysis does not include individuals from the BAGS cohort, since they alone were sequenced directly by Illumina Inc. This within-sequence-center analysis also revealed significantly fewer DNMs per individual in the Amish compared with European individuals from FHS (Anova, P<0.005), whereas we found no significant difference in DNMs per individual between European American individuals from CFS, African American individuals from CFS, and Latino individuals from CRA (Figure S9B,S9C). These results persist after adjusting for parental age, (Anova, P<1.76×10^-3^, Figure S9D), and suggest that differences between ancestry thought to influence the DNM rate, such as heterozygosity, do not appreciably shape the accumulation of *de novo* mutations. When repeating these analyses after randomly choosing one offspring per nuclear family, our results are qualitatively the same. This was the case for all of our analyses that focus on per individual DNM measures, and for simplicity, we only report results from analyses run with our full dataset (except when noted with our heritability models).

**Figure 3.**
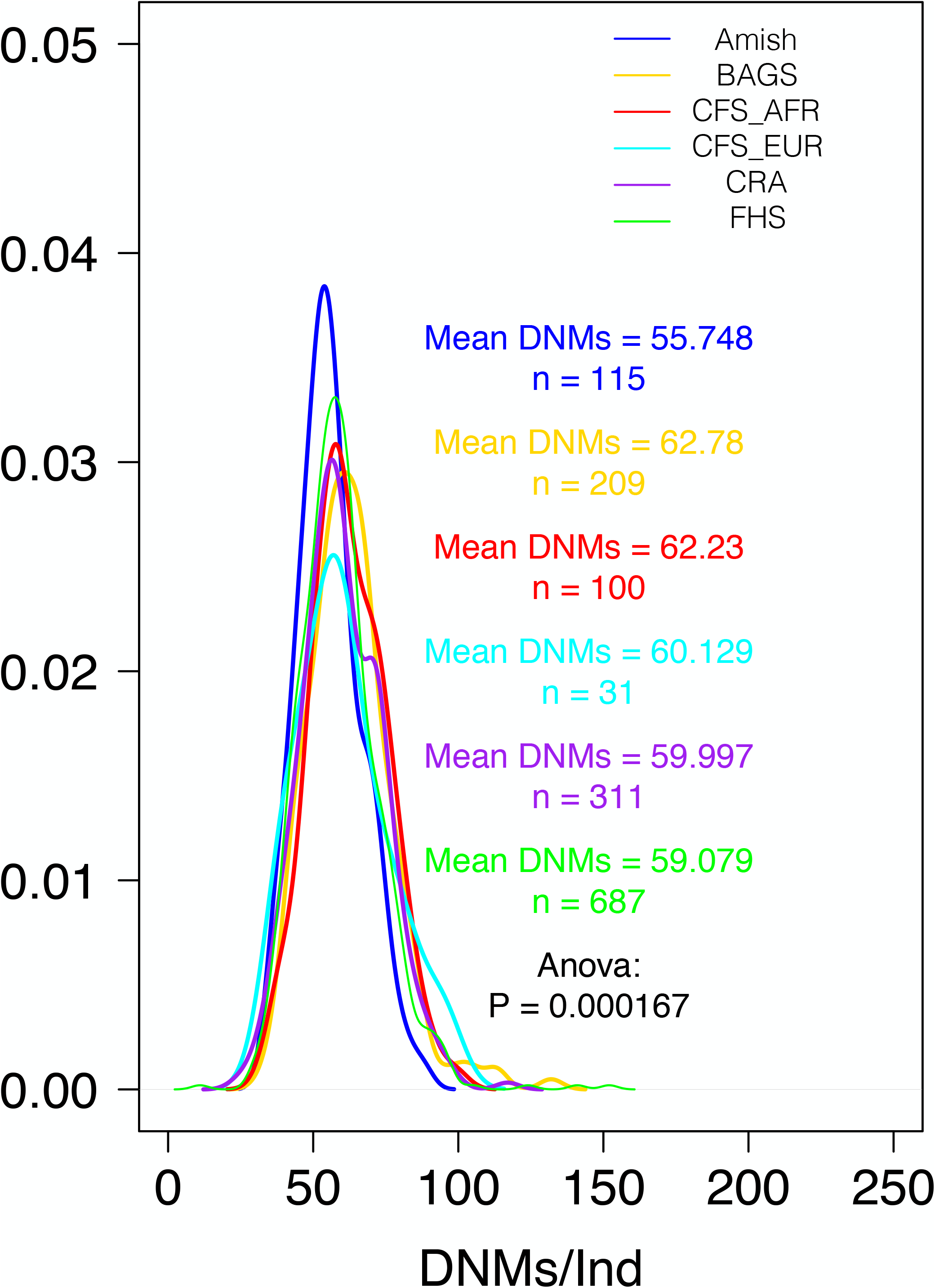
SNV DNM totals across diverse cohorts. SNV DNM totals per individual show significant differences across cohort, which seem to be driven by a reduction in the Amish.

To test more directly how inter-ancestral differences might influence DNM accumulation, we assessed the relationships between DNM count and African ancestry proportion, and DNM count and Native American ancestry proportion. Within the BAGS, CFS_AFR, and CRA individuals, which represent the three cohorts with significant African ancestry, we found no significant correlation between African ancestry proportion and DNM count after adjusting for parental age (P=0.647, Figure S10). This absence of a relationship persists when looking only within BAGS individuals or only within CRA individuals (Figure S11A, S11B), and there is also no relationship when comparing Native American ancestry proportion and DNM count within Latino ancestry individuals from the CRA cohort (Figure S11C).

When evaluating whether single base or 3-mer mutation frequencies correlate with global genomic ancestral proportion, we do not find significant differences. However, TCC→TTC 3-mer mutations showed a trend of being negatively associated (P<6.5×10^-3^) with European ancestry proportion within the BAGS cohort of admixed African ancestry individuals (Figure S12). This is interesting when considered in conjunction with recent evidence that an ancestry specific pulse increased the occurrence of these 3-mer mutations in Europeans^35^. We also find significant differences in DNM totals across families with ≥ 2 children (P<0.005) (Figure S13).

### Shifts in the mutational spectrum of the Amish founder population

To further assess the reduced number of SNV DNMs seen in the Amish, we compared DNM counts for each single base mutation type across cohort. Linear regression reveals differences in C→A, T→C, and T→G mutation across cohort, which are largely driven by decreases in these mutations in the Amish and increases in these mutations in individuals from the BAGS cohort (Figure S14). To further evaluate these differences while controlling for sequencing center, we compare mutations between the Amish and FHS cohorts, and find the reduced number of T→C mutations in the Amish to persist in this conservative sub analysis (Figure 4). C→A mutations also show marginally significant reductions in the Amish compared to FHS (P=0.053), although the reduction in T→C mutations is notably more significant (P<8×10^-4^), and explains about 50% of the overall DNM reduction seen in the Amish.

**Figure 4.**
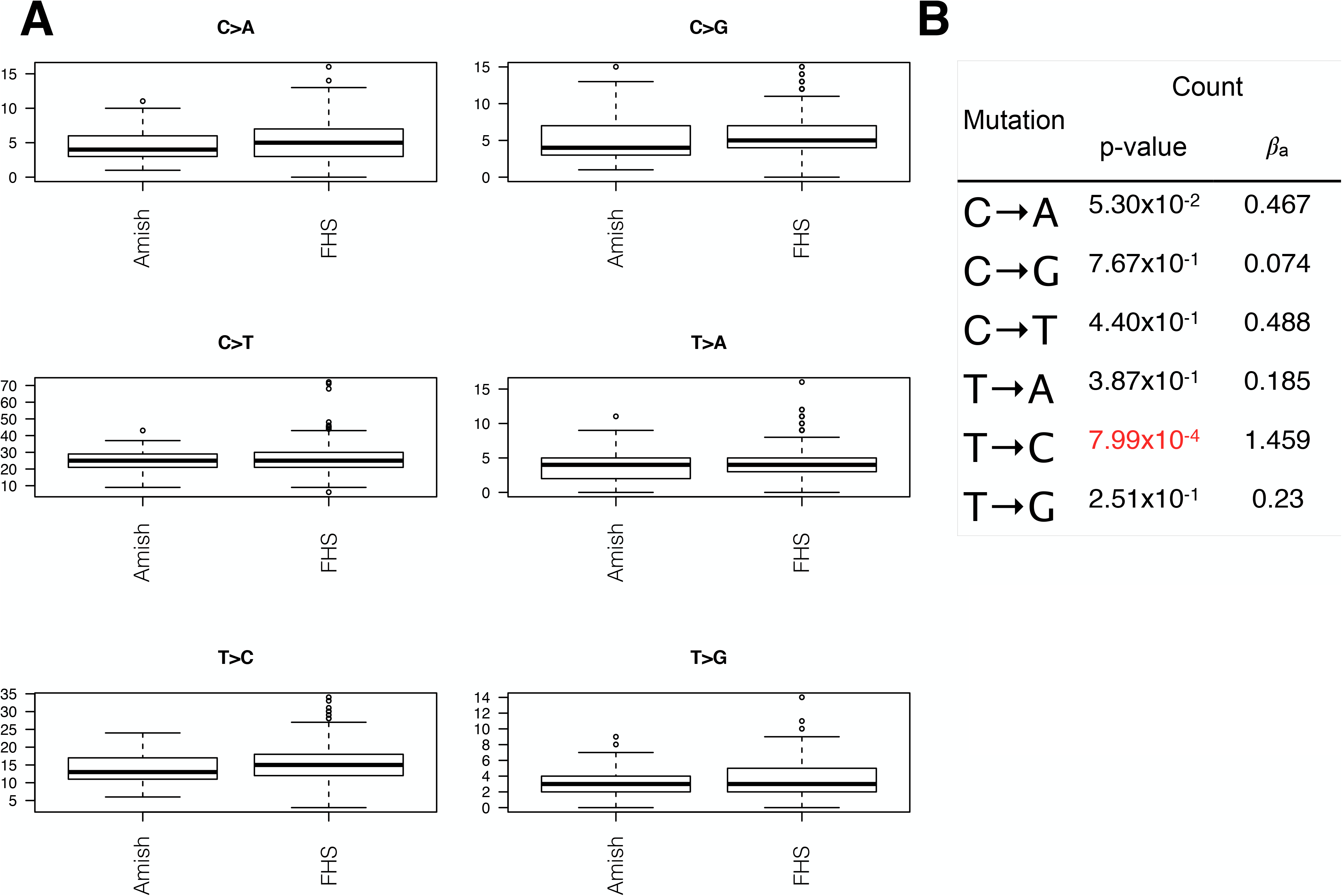
Single base mutation differences between Amish and FHS. Amish individuals have significantly fewer T→C mutations, which explain about 50% of their overall SNV DNM reduction (A and B). C→A mutations show a marginally significant reduction in the Amish compared with FHS.

### Local ancestry analysis does not identify ancestrally distinct mutation rates

To look more closely at whether mutation rates differ between ancestries, we compare the rates of DNM accumulation across ancestrally distinct genomic segments within admixed samples. Using local ancestry assignments (see Methods), we count DNMs across all possible diploid ancestral segment combinations (e.g. homozygous African, heterozygous African and European, etc) (Figure S15), and limit each comparison to individuals with at least 80 megabases of each diploid category. The resulting intra-individual comparisons allow for the control of unmeasured variables that may otherwise confound inter-individual analyses (for example environmental exposures that increase mutation rate), since the only variable that should differ between diploid segments is their ancestral origin, and any effects of this ancestral background on local sequence context. After estimating DNM rates per individual by normalizing DNM counts by diploid category base total (i.e. rates per base pair), we do not find evidence of ancestry specific differences in DNM rate (Figure S16). While we did find a significant reduction in the mutation rate in African/European segments compared with African/African segments (P<6.1×10^-4^), other comparisons between African and European segments show no differences. Given the absence of consistent differences in SNV DNM rate across local ancestral segments, these results do not provide compelling support for the existence of ancestry-based differences in SNV DNM rate.

### The heritability of de novo mutation accumulation

To assess whether we could use genome-wide association analysis (GWAS) to detect any genetic loci that might underpin the inter-individual variation we see in DNM accumulation, we first wanted to test for what proportion of the variation in DNM accumulation was explained by genetics. One measure of this is narrow-sense heritability (h^2^), which represents the proportion of phenotypic variation explained by additive genetic effects^42^, and which can be used as a null model for running associations at every locus (i.e. each locus is tested for the proportion of h^2^ that it explains). Treating the number of SNV DNMs per individual as a phenotype reflecting DNM accumulation, we used the MMAP software^43^ to estimate the h^2^ of DNM accumulation with a variety of models (see methods). When using all samples across all cohorts, we used father’s age at offspring’s conception, mother’s age at offspring’s conception, and cohort label as covariates, and a genetic relatedness matrix (GRM) to account for sample relatedness. Very interestingly, we estimate an h^2^ = 0.0 (SE = 0.028) with this analysis, and we reaffirm a father’s age effect (P<2.5×10^-59^), a mother’s age effect (P<6.96×10^-7^), and an effect of Amish cohort status (P<2.99×10^-4^) (Table 2, S3). When running this base model on each cohort separately, h^2^ estimates are zero or near-zero (i.e. non-significantly above zero) for each cohort (the CRA model didn’t converge, but had at least one optimum that estimated h^2^ at zero).

**Table 2.** Heritability Model across all cohorts. Results are shown from heritability models run with MMAP across all samples with paternal and maternal ages available (n=1388). Heritability is estimated as zero (h^2^ = 0.00), with a standard error of 0.03). These models confirm that paternal age at offspring’s conception (P = 4.61×10^−65^), maternal age at offspring’s conception (P = 2.14×10^−8^), and Amish cohort status (P = 1.09×10^−4^) are significantly correlated with DNM total per individual.

| Variable | Value | Variable | Value |
| --- | --- | --- | --- |
| Mean DNMs | 59.57 | βpaternal_age Standard Error | 0.07 |
| SD | 13.38 | βpaternal_age P-value | 4.61E-65 |
| Min | 11.00 | βmaternal_age | 0.43 |
| Max | 124.00 | βmaternal_age Standard Error | 0.08 |
| Kurtosis | 0.74 | βmaternal_age P-Value | 2.14E-08 |
| DNMs > 3 SD | 6.00 | βAmish | -5.18 |
| DNMs < 3 SD | 1.00 | βAmish Standard Error | 1.33 |
| Sample Size | 1388 | βAmish P-value | 1.09E-04 |
| Number of Covariates | 8.00 | βBAGS | -0.90 |
| h2 | 0.00 | βBAGS Standard Error | 1.27 |
| h2 P-value | 0.00 | βBAGS P-value | 0.48 |
| h2 P-value Standard Error | 0.03 | βCFS_AFR | 0.63 |
| Proportion Variance Explained By Covariates | 0.51 | βCFS_AFR Standard Error | 2.15 |
| Adjusted Proportion Variance Explained By Covariates | 0.51 | βCFS_AFR P-value | 0.77 |
| In Likelihood | -3843.13 | βCRA | -1.37 |
| Likelihood Method | MLE | βCRA Standard Error | 1.13 |
| Intercept | 13.00 | βCRA P-value | 0.22 |
| Intercent Standard Error | 1.76 | βFHS | -2.56 |
| Intercept P-value | 2.46E-13 | βFHS Standard Error | 1.05 |
| βpaternal_age | 1.24 | βFHS P-value | 0.01 |

To evaluate whether the portion of mutational variation explained by parental age might be shaped by additive genetic factors, we repeated these heritability models without adjusting for either parent’s age at conception. Interestingly, these models estimate non-zero heritability (h^2^ = 0.239, P<2.95×10^-4^), with the BAGS cohort contributing most significantly to this non-zero heritability estimate. However, when repeating all analyses with only one offspring randomly chosen per nuclear family (i.e. removing sibling data), h^2^ is zero in all models, suggesting that the significant non-zero heritability we see here when not adjusting for parental age is driven by the shared parental contribution of siblings and not by genotypic similarity. This is consistent with both the significant inter-family variation and the lack of inter-ancestral variation we see when analyzing SNV DNM rates.

## Discussion

Here we call *de novo* mutations across 1,465 individuals from diverse cohorts sequenced through the TOPMed program. Our call set is determined by a filtering heuristic that is similar to previous approaches^19^, and its specificity is supported by a Bayesian approach implemented in the TrioDeNovo software^44^. Furthermore, the significant correlations we find between DNMs and paternal and maternal age at conception, and the consistency of our mutation rate estimates with previous genetic estimates, lend additional support for the veracity of our call set. We confirm that C→T and T→C transitions are the most frequent mutation type among DNMs, and demonstrate that CpG mutations in a 3-mer context are highly overrepresented after accounting for background 3-mer frequency.

Despite this predominance of C→T mutations amongst DNMs, we do not find any differences between populations or ancestral backgrounds in C→T mutations. This is interesting given the high frequency of this mutational class, and it suggests that processes driving cytosine deamination are fairly conserved. This is consistent with the assertion by others that CpG mutation rates may serve as a better “molecular clock” than the base substitution rates that have previously been used^17,34^. Similarly, we do not find differences in other single base and 3-mer mutation types across ancestral background. While some of this might relate to having limited statistical power, it suggests that mutational processes and accumulation are quite similar between ancestral groups. However, we do find the TCC→TTC mutation to be the most significantly negatively associated with European ancestry proportion within the BAGS cohort, which has the highest average African ancestry proportion among our cohorts. While this correlation was nonsignificant after multiple testing correction (P<6.4×10^-3^), this result is interesting when contrasted with recent findings of Europeans having had a significantly higher rate of TCC→TTC mutations in the past^34,35^. Whether there are in fact no current differences between ancestral background in the rate of this mutation, or whether Africans possibly have a higher current rate than Europeans, our result is consistent with these earlier findings in suggesting that the rate of this mutation evolves quickly. In other words, assuming the mutation rate of the TCC→TTC 3-mer experienced a short population-specific increase, our results suggest it subsequently experienced a rate decrease in European ancestry individuals and/or a rate increase in African ancestry individuals, which provides support for the rapid evolution of mutational processes.

The correlations we find between local recombination and DNM rates are concordant with research showing that processes underpinning recombination are mutagenic^25,26,30^, and our model comparing population-level rare variation with both recombination and DNM rates adds further resolution to the relationship between recombination and mutation. Areas of the genome with the highest local DNM rate are in regions with high recombination that have recently been identified as featuring specific recombination-based processes that drive mutation^24^. These processes seem to underpin specific mutational patterns in maternal gametes, and to preferentially drive C→G mutations. In accordance with this, we find these regions to have increased proportions of C→G mutations, and we find high recombination regions in general to have twice the C→G frequency as low recombination regions. Furthermore, we find genome-wide correlations between mutation type and local recombination rate, with C→T and C→G mutations correlating positively with local recombination rate and T→C and T→G mutations correlating negatively (Figure S3). Alternatively, regions with high recombination rate estimates that have low DNM rates (Figure 2, tall blue bars) are potentially good candidates for the identification of features that may explain less mutagenesis than recombination rate alone would predict, such as early replication times^30,45^, or of circumstances in which recombination itself may be less mutagenic. While recent research into the relationship between DNMs and fine-scale local recombination rates identified up to 50-fold increases in DNM rates around meiotic crossover events^30^, the fact that these events are rare is likely why we only explain about 3% of the variation in DNMs on the basis of recombination. This recombination study also found 35 loci influencing recombination rate, which may lead one to ask why we don’t find any signal of genetic influence over DNM accumulation (i.e. heritability > zero) given that recombination is mutagenic and associates with DNMs in our dataset. First, this is likely due to the fact that recombination only explains a small portion of the overall variation in DNMs, and small effect size modulators of recombination are unlikely to be detected as correlates of DNM rate without enormous sample sizes. Second, we adjust for parental age effects, which would eliminate from many of our heritability models the portion of variation in DNM rate influenced by meiotic recombination. Overall, these results help to inform questions about where in the genome these relationships are strongest.

This model also estimates the quantitative contribution of mutation to population level variation. Surprisingly, we are able to explain up to 35% of the variation across the genome in population level rare variation using local recombination and DNM rates, with nearly 31% of this attributed to local DNM rates. Regions with high mutation rates have the most variation in the genome (e.g. chr8, chr16), and also have high recombination rates that likely increase mutation accumulation. Numerous recent studies have found complimentary results that describe the types of mutations predominating in high variation regions and the likely sources of these mutations^21,22,24^. While some of this recent work demonstrated that clustered DNMs contribute very significantly to clustered SNPs (specifically C→G SNPs), clustered DNMs make up less than 2.5% of DNMs^24^. Here we show that SNV DNMs contribute profoundly to the entirety of rare single-nucleotide variation, and our integrative analysis provides additional insight into how mutation shapes genetic variation across the genome.

While the Heterozygosity Instability hypothesis predicts that increasingly heterozygous genomes will have higher mutation rates^31,32^, we only find evidence of a weak relationship between heterozygosity and DNM rate. Furthermore, we don’t find the differences in DNM rate between ancestral background that differences in heterozygosity across ancestry would have predicted.However, very interestingly, we do find a reduced mutation rate in the Amish, which seems to be driven by a reduction in T→C mutations, and which does not seem to be underpinned by genetics. Despite being a founder population that diverged from other Europeans only very recently^46^, the Amish show a mutation rate reduction of about 6%, which persists when the Amish are compared with other European ancestry individuals from the Framingham Heart Study (FHS) in a sub-analysis that controls for sequencing center. Together with our estimation that SNV DNMs have zero narrow-sense heritability, this suggests that the environment may play a bigger role in modulating the mutation rate than previously appreciated. The Amish lifestyle features pre-industrial era aspects, and while modern Amish communities are diverse and have adapted to the usage of some modern items, they continue to limit the influence of technology in their daily lives^47,48^. Given this, it is possible that the Amish are exposed to fewer mutagens, and that this “clean living” may be partially responsible for the reduced mutation rate we report here. For example, studies have shown that rural areas, such as those similar to the areas occupied by the Amish, have fewer carcinogens and mutagens than industrialized areas^49-51^. Polycyclic Aromatic Hydrocarbons (PAHs) in particular represent a major pollutant and carcinogenic mutagen^52^ that the Amish are likely to be significantly less exposed to due to their rural and minimally industrialized lifestyles^49-51,53,54^. Given that PAHs and other mutagens have been found to correlate with traffic levels^54-57^, to contaminate food from industrialized regions, and to be increased by tobacco usage^49^, and that the Amish eschew driving in favor of horse and buggy, feature a locally grown organic diet rich in vegetables, entirely avoid tobacco usage, and maintain physically active lifestyles^47,48^, there is at least significant rational for reduced mutagenesis in the Amish^49,58,59^. Furthermore, the specific mutations we find most reduced in the Amish (i.e. T→C, T→G, and C→A) show the largest increases in experiments assessing the mutagenic effects of oxidative damage on DNA repair^60^. One of the most common oxidatively modified DNA bases (e.g. 8-OH-dGuo) drives C→A mutations, and another common oxidative lesion (e.g. thymine glycol) increases the rate of T→C transitions^60^. Recent analysis of mutation patterns has also called into question the canonical view that DNMs rise predominantly from replicative errors, and suggests that exogenous mutagens may play a larger role in mutation accumulation than previously appreciated^23^. If the Amish do in fact experience less environmentally driven mutagenesis, then one would predict a significant reduction in the rate of cancer in the Amish. This is exactly what has been found in multiple Old Order Amish populations, with a particularly large reduction in cancer rates found in men^61,62^. A similar reduction in overall mortality has been found in Amish men compared with FHS men, which is hypothesized to be due to lifestyle factors such as reduced tobacco use and increased physical activity^47^. Given that DNM mutation in sperm is the single largest driver of DNM accumulation, an Amish environment that potentially limits DNA damage in Amish men is consistent with a reduction in DNM rate. In fact, while not significantly different than the paternal age effects found in the other cohorts, the number of DNMs attributable to each year of paternal age at offspring’s conception is lowest in the Amish (Table S4).

When viewed in conjunction with our finding that variation in DNM accumulation does not appear to be heritable, the idea that the environment may explain the reduction in DNMs found in the Amish becomes increasingly plausible. While we acknowledge these potential explanations as entirely speculative, our findings along with the aforementioned context suggest environmental influence as a plausible explanation worthy of additional consideration and follow-up. The fact that the only signal of DNM heritability we detect is driven by siblings when not accounting for parental age effects further suggests that the environmental similarity shared by siblings (including parental age effects) is significantly influencing DNM rate. In sum, the mutational differences we find in the Amish stand in contrast to the homogeneity seen across the other diverse human populations we analyze, and suggest that additional work is needed to better appreciate the forces shaping human mutational processes at fine-scales.

## Acknowledgements

Whole genome sequencing (WGS) for the Trans-Omics in Precision Medicine (TOPMed) program was supported by the National Heart, Lung and Blood Institute (NHLBI). WGS for “NHLBI TOPMed: Whole Genome Sequencing and Related Phenotypes in the Framingham Heart Study” (phs000974) and “NHLBI TOPMed: Genetics of Cardiometabolic Health in the Amish” (phs000956) were performed at the Broad Institute of MIT and Harvard (3R01HL092577-06S1, 3R01HL121007-01S1). WGS for “NHLBI TOPMed: The Genetics and Epidemiology of Asthma in Barbados” (phs001143) was performed by Illumina, Inc (3R01HL104608-04S1). WGS for “NHLBI TOPMed: The Cleveland Family Study (WGS)” (phs000954) and “NHLBI TOPMed: The Genetic Epidemiology of Asthma in Costa Rica” (phs000988) were performed at the University of Washington Northwest Genomics Center (3R01HL098433-05S1, 3R37HL066289-13S1). Centralized read mapping and genotype calling, along with variant quality metrics and filtering were provided by the TOPMed Informatics Research Center (3R01HL-117626-02S1). Phenotype harmonization, data management, sample-identity QC, and general study coordination, were provided by the TOPMed Data Coordinating Center (3R01HL-120393-02S1). We gratefully acknowledge the studies and participants who provided biological samples and data for TOPMed.

## Author Contributions

MDK and TDO designed all experiments and wrote the manuscript. MDK performed all bioinformatic analyses under the guidance of TDO, and with analytic input from JAP, JRO, BDM, SRB, SZ, DAN, and RH. DLP ran heritability models with input from MDK, JAP, JRO, and TDO. NLHC, BEC, HW, MD, JZ, MES-Q, LA, and JCC collected samples and /or provided meta data as part of a participating TOPMed cohort. STW, KB, SSR, RSV, ADJ, RAM, JGW, DAN, GA, JRO, JAP, and BDM coordinated sequencing and/or helped with project design as a TOPMed or cohort investigator. All authors were involved with editing the manuscript.

## Methods

*TOPMed DNA samples, sequencing, and data processing*

Whole genome sequencing (WGS) was performed using whole blood from samples previously collected and consented across 90 NHLBI funded research projects (7 Framingham samples were sequences using cell lines as opposed to blood^37^). Samples were sequenced using paired end approaches using Illumina HiSeq X Ten machines, and featured 2 × 150 bp reads and a mean depth of 38X. All sequencing followed the protocol in the Illumina TruSeq PCR-Free Sample Preparation Guide (Illumina cat# FC-121-2001), and used PCR-free library preparation kits purchased from KAPA Biosystems (see https://www.nhlbiwgs.org/topmed-whole-genome-sequencing-project-freeze-5b-phases-1-and-2 for additional details). Periodic sequence data processing and analysis was performed to produce genotype data “freezes” that included all the available samples at the time of the release. Downstream alignment and processing was done according to a published protocol^63^ (also see https://github.com/CCDG/Pipeline-Standardization/blob/master/PipelineStandard.md), although the data freeze we used aligned sequences to the build 37 of the human genome reference using bwamem^64^. Joint variant calling was then done using the GotCloud13 pipeline for all samples available in a given freeze. This generated a single, multi-study, genotype call set. A support vector machine (SVM) variant site quality filter was trained using known variants and Mendelian-inconsistent variants as positive and negative training sets, respectively (see https://github.com/statgen/topmed_variant_calling for more details). To improve quality control at the level of the sample, checking was done for pedigree errors, discrepancies between self-reported and genetic sex, and concordance with prior genotype data. Details regarding WGS data acquisition, processing, and quality control are described on the TOPMed website^65^, and are further documented in each dbGaP release accession.

### TOPMed cohorts and samples

This study utilizes samples from six cohorts from the Trans-Omics for Precision Medicine (TOPmed) program. These consist of Amish individuals from the Genetics of Cardiometabolic Health in the Amish study (Amish), Barbadians from the Barbados Asthma Genetics Study (BAGS), European and African Americans from the Cleveland Family Study (CFS_EUR and CFS_AFR), Costa Ricans from the Genetic Epidemiology of Asthma in Costa Rica (GACRS) and Childhood Asthma Management Program (CAMP) studies (collectively referred to as CRA), and European Americans from the Framingham Heart Study (FHS). As detailed in Supplementary Table 1, we utilized samples from a total of 1,465 children across these cohorts, which came from a total of 1,214 independent families. Including parents, this study used whole genome sequencing from 3,893 samples. After calling DNMs, we removed a total of 12 outlier samples from any follow-up analysis: six were extreme outliers, and were determined to derive from pedigree errors (i.e. incorrect parentage within our meta data), while the other six were outliers beyond five standard deviations, which likely resulted from issues with sequence quality. This left us with 1,453 samples for follow-up analysis. Repeated analyses restricting to one randomly chosen offspring per family used 1,205 samples after the removal of outliers. We also had a variety of meta data for the majority of samples, including fathers age at offspring’s conception, mothers age at offspring’s conception, sex, birth age, DNA age, and sequencing center.

### Filter approach for calling DNMs

In order to identify candidate *de novo* mutations with a low false positive rate, we implemented a conservative heuristic approach that relied on stringent filtering. We applied this approach to all trios (two parents and one offspring) from six TOPMED cohorts (Amish, BAGS, CFS_EUR, CFS_AFR, CRA, FHS) for whom we had genotype information for each trio individual. Since these cohorts were selected on the basis of their enrichment for family data, we had data for multiple children in numerous families. For these families, each offspring plus parent trio was considered independently, and we repeated all analyses with only one randomly chosen offspring per family in order to ensure that our conclusions are robust to family structure. Beginning with variant call format (VCF) files containing genotypes called by the TOPMed consortium data processing team for all 18,526 individuals in freeze 4, we assessed all variants in all trio individuals from 1465 trios. After excluding all variants that did not pass best practices quality control (represented by the lack of a “PASS” in the VCF file filter column), we further evaluated all variants that were heterozygous in the offspring and homozygous reference in each parent. Remaining variants were excluded if they had total allele depth less than ten reads (AD < 10), genotype quality less than thirty (GQ < 30), a heterozygous to homozygous reference genotype likelihood ratio less than 1×10^10^, a ratio of reference allele read count to alternative allele read count less than 0.3 or greater than 0.7 (reference reads/alternative reads < 0.3 or reference reads/ alternative reads > 0.7), total paternal allele depth less than 10 reads, total maternal allele depth less than 10 reads, paternal genotype quality less than 30, maternal genotype quality less than 30, or greater than or equal to 1 alternative allele read in either parent (paternal_alt_reads > 0 or maternal_alt_reads > 0). After the exclusion of 12 samples, 6 of whom involved pedigree errors due to incorrect parentage, and 6 of whom were outliers beyond 5 standard deviations, this filtering-based heuristic resulted in the identification of 86,865 de novo mutations (DNM) across 1453 samples.

### Alternative DNM calls using Triodenovo software

To provide further support for the accuracy of our SNV DNM call set (i.e. that it has a very low false positive rate), we used the Bayesian *de novo* mutation caller called TrioDeNovo^44^ to call SNV DNMs across 5 autosomes (chromosomes 18-22, representing ∼10% of the human genome) in all 1,465 samples. All SNV DNMs identified across these autosomes in all samples by our filtering approach were identified in the TrioDeNovo call set, which provides additional support for the veracity of our filtering approach call set. Interestingly, TrioDeNovo also identified ∼2.5 times as many SNV DNMs per individual as our filtering approach, which further supports the usage of our strict filtering for the elimination of the false positives that other approaches tend to identify.

### Mutation rate Estimation

Given 0.9916 as the proportion of the genome covered by a read depth of 10+ across freeze 4 of the TOPMed sequencing project, and 0.9875 as the mapping proportion, we calculate 3.0 × 10^9^ * 0.9916 * 0.9875 = 2.938 x10^9^ as the number of accessible bases for this dataset. Using 86865/1453 = 59.78 as the estimated number of DNMs per individual, we then calculate a mutation rate estimate of as 59.78 DNMs / (2.938 x10^9^*2 accessible bases per diploid genome) = 1.017 × 10^-8^.

### Analyzing single base and 3-mer mutation types

For each of the 86,865 single nucleotide DNMs in our call set (excluding outlier samples), single base type was determined by the base change for each DNM, with mutation types related by reverse complementarity totaled together (e.g. T→G mutations represent T→G + A→C). Across this entire call set, proportion of each type was calculated as single-base mutation type count divided by total SNV DNMs in the call set. Associations with parental ages at offspring’s conception were determined using a generalized linear model with Poisson outcomes described by the following equation:

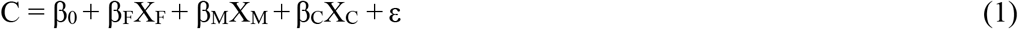

where C represents the total count for each single base mutation across the entire SNV DNM call set, X_F_ is the paternal age at offspring’s conception, and X_M_ is the maternal age at offspring’s conception. The distribution of single base mutation types across the genome was generated by calculating the proportion for each possible single base mutation across one megabase (Mb) windows genome-wide. When visualizing these distributions, we excluded the final segment from each chromosome, which was typically shorter than one Mb, as well as windows with fewer than 5 DNMs or a recombination rate estimate of zero (as this was highly correlated with the presence of centromeres). 3-mer DNMs were annotated by evaluating the reference base preceding and following each DNM, and counts for each of 96 possible 3-mers (after collapsing complimentary 3-mers together) were totaled across the entire SNV DNM call set. We also calculated the frequency of each 3-mer in the GCRh37 reference genome, and then normalized each SNV DNM 3-mer count by its frequency in the reference genome. Single base mutations and 3-mers were also totaled per sample, and linear models (Poisson and normal) were run with father’s and mother’s ages at conception of the offspring as covariates in order to test the relationship between global ancestry (European, African, Native American) and single base or 3-mer mutation count.

### Correlations with recombination and rare variation

To estimate the relationship between mutation rate and population-level rare variation, and between mutation rate and recombination, we calculated local SNV DNM rates, local recombination rates, and rare variation levels for one Mb windows across the autosomes. SNV DNM rates were estimated using the number of SNV DNMs from our 86,865 DNM call set that overlapped each 1 Mb window, and were scaled as the local mutation rate per person per base according to the following equation:

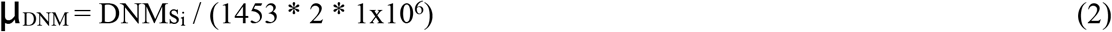

where μ_DNM_ is the local mutation rate, DNMs_i_ is the DNM count in window i, 1453 represents the number of samples from which our SNV DNM call set is derived, and 1×10^6^ scales per base pair. To calculate local recombination estimates across each 1Mb window, we used genetic map data from HapMap (cite), and used GrCh37 or GRCh38 annotations depending on the analyses (see below). Local recombination rate was estimated as the first genetic position estimate in a window minus the last, divided by the total number of bases separating these estimates, and scaled to 1 million. The result of this calculation was in units centimorgans per megabase (cM/ Mb), and was used as the recombination rate estimate for the 1 Mb window. Using WGS data from ∼41k diverse and unrelated samples from freeze5 of the TOPMed consortium^37^, we calculate the total number of variants with allele frequencies ≤ 0.005 in each 1Mb window, normalize these counts as Z-scores, and use these Z-scores as measures of the levels of recent variation existing in the human population across the genome. Given that most variation is rare, and that rare and common variation are highly correlated^37^, these rare variant based estimates also reflect how overall genomic variation varies across the genome. To control for the potential effects of coding variation on rare variant totals, as well as for the potential effects that regions with mappability concerns may have on both rare variation and DNM calling (in ways that could potentially be confounded), we used linear regression to adjusted TOPMed rare variation levels by the proportion of bases that are coding and the proportion of bases that were identified in TOPMed as having mappability concerns. The residuals from these regression analyses were used as adjusted levels of rare variation in a repeat analysis of the relationship between DNM rate, rare variation, and recombination rate. Given that this TOPMed data is annotated using the GRCh38 version of the human reference genome, all data included in analyses using these TOPMed rare variation measures were annotated using the GRCh38 human reference.Specifically, when appropriate, we used genetic map data from GRC38 to calculate local recombination rates, and we converted the positions of our SNV DNMs from GRCh37 to GRCh38 with UCSC’s liftOver tool^66^ before estimating local mutation rates. Otherwise, analyses including local recombination and DNM rates used data based on GRCh37 annotations. To assess the relationship between TOPMed variation levels, recombination, and mutation, we used linear regression. By using both local recombination rate and local SNV DNM rate as independent variables, and TOPMed rare variation transformed into Z-scores as our dependent variable, we estimate how much variation in population level genomic variation is explained by both recombination and mutation. To see how much of this variation is explained by mutation after accounting for recombination, we first regressed TOPMed variation onto recombination and then regressed the residuals of this model onto mutation.

For comparisons of mutation proportions across windows, single base SNV DNM totals were calculated and converted to proportions by dividing by the total number of DNMs per 1Mb window. High variation, high DNM, high recombination 1Mb regions, as seen in Table S2, are defined as having rare variant Z-scores in the top 10% of values, and DNM and recombination rates in the top 20%. High recombination regions are those with recombination rate values in the top 20%, and low recombination regions are those with Z-scores ≥ −2, DNM rates in the top 80% of values, and recombination rates in the bottom 20% of values. To test the relationship between recombination and mutations, we regressed local recombination rate onto local SNV DNM rate, and repeated similar analyses independently per chromosome to see how the relationship between recombination and mutations varies across the autosomes.

### Correlations with Heterozygosity

To test whether more heterozygous genomes have a higher DNM rate, we calculated genome-wide heterozygosity scores for each of the 1,453 samples included in our analysis. These were compared with the total SNV DNM count per individual using linear regression, and for comparison within only a particular ancestry, samples were subset down accordingly. To account for the effects of parental age on DNM count, we regressed out paternal and maternal age at conception, and then reran regression models between this adjusted DNM value and heterozygosity.

### Local ancestry analysis

We phased the data using the Eagle2 algorithm as implemented in the Eagle software^67^. For this phasing, we used binary input files, GRCh37 genetic map files, 4 threads, and the --maxMissingPerIndiv 0.6 argument. Next, we combined whole genome sequencing (WGS) data from the 1000 genomes with high coverage WGS data from the Peruvian Genome Project^68^ to assemble genotype data from 1,752 samples of European (499), African (640), East Asian (498), and Peruvian (115) ancestry. This genotype data was used as reference input to the RFMIX software^69^, which we used to estimate local ancestry across all 1,465 of our samples for which we called DNMs. Using these local ancestry calls, we went through the genomes of each sample (1453 after removing outliers, see *TOPMed cohorts and samples*) and binned contiguous regions by their ancestral origins. In other words, a region was called European-European if our local ancestry calls estimated both chromosomal copies of that region as having European ancestral origin. Once every region in the genome had been annotated in this manner with diploid categorization, the DNMs in each region were tallied for each individual. Finally, to calculate local estimates of *de novo* mutation rate across these ancestrally distinct diplotypic segments, the DNM counts and base totals for each diploid region within each individual were summed, and the resulting DNM count totals were divided by the total number of bases, per diploid region. Within each diploid comparison (i.e. European-European vs European-African), we excluded individuals who did not have at least 80 megabases of each diploid category (minimum 2 expected DNMs) in order to avoid the noise that arises from working with exceedingly small numbers. The resulting distributions were compared using an analysis of variance.

### Mutation Spectrum analysis

Using the single base changes we previously determined, we counted the number of each mutation type per individual for each of the 1,453 samples included in our overall analyses. After using regressing to adjust for both paternal and maternal age at conception, we used logistic regression to test for a relationship between mutation type count and population label.

### Heritability analysis

We treated the number of SNV *de novo* mutations per individual as a quantitative phenotype, and estimated its heritability with the MMAP software^43^, which estimates narrow-sense heritability (h^2^) using a linear mixed model. The following equation represents our base model, which we adjust for subsequent analytic iterations, as described below:

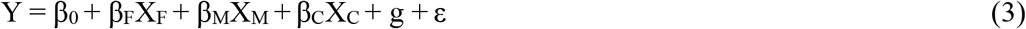

where X_F_ is the paternal age at offspring’s conception, X_M_ is the maternal age at offspring’s conception, X_C_ is a categorical variable indicating study, and g is the random effects, where Var(g) = Σ^2^_g_ * A, and A is the genetic relationship matrix (GRM). We used this base model to estimate h^2^ across a total of eight separate analyses, with all six cohorts together serving as a sample set, each separate cohort serving as a sample set, and the Amish and FHS cohorts serving as a single pooled sample set. In all of these association analyses, we remove samples (n=3) with SNV DNM values ≥ 130 in order to more stringently remove outliers and reduce the likelihood of spurious associations. For the all cohort analysis, cohort membership was included as a covariate. In order to evaluate whether the portion of variation in SNV DNM accumulation attributable to parental age was itself heritable, we repeated the above models without adjusting for parental ages as covariates. These base models are run in MMAP with the following sample command and parameters (which change depending on covariates etc):

*mmap --ped $ped --read_binary_covariance_file $grm --phenotype_filename $pheno--phenotype_id PROBAND_ID --num_mkl_threads 4 --model add --trait $trait --file_suffix $suff --single_pedigree --covariates $covariates*

where “$ped” is a pedigree file containing family IDs, parent IDs, and sex, “$grm” is a genetic relationship matrix, “$pheno” is a phenotype file containing DNM values and relevant covariates,“--phenotype_id” specifies which column contains the subject ID in the phenotype file, “$trait” specifies the column name of the phenotypic trait to utilize in the model, “--model add” specifies the use of an additive model,“$suff” specifies a file suffix to prevent overwriting, “-num_mkl_threads” specifies the number of threads used, “--single-pedigree” indicates that the family ID in the pedigree file will be ignored, and “$covariates” specifies the column names of the covariates to be included in the model from the phenotype file. When ignoring the family ID and specifying a GRM, the family structure is determined via the GRM rather than the pedigree file. Heritability calculations in MMAP use the GRM alone as its genotypic informational input. Overall, this command enables the estimation of heritability (h^2^), specifies a GRM, and fits an additive linear mixed model. Heritability analyses that adjusted for parental age effects produced qualitatively similar results when repeated with one randomly chosen offspring per nuclear family.

---

**Figure S1.**
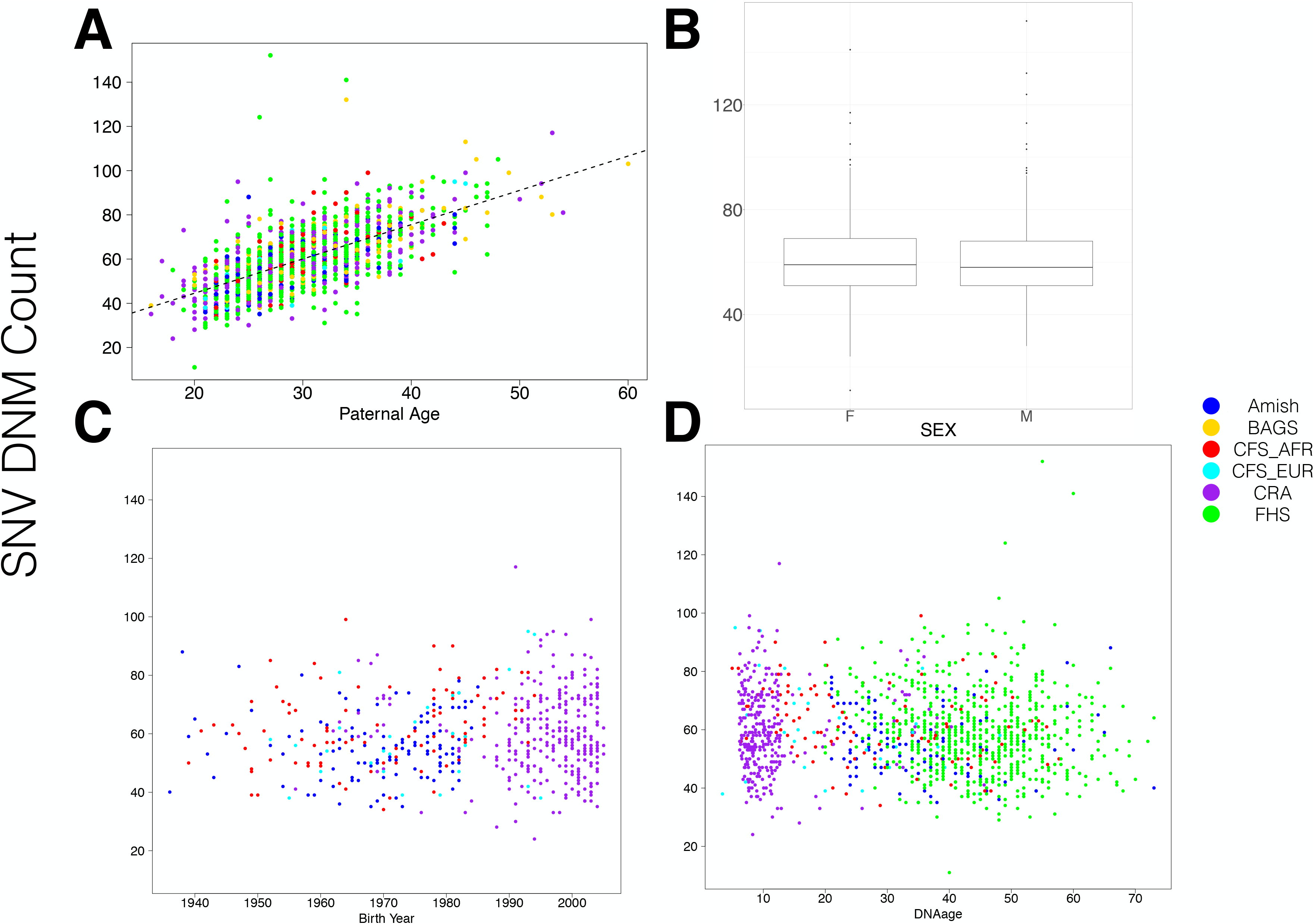
Correlations between SNV DNM totals and various meta data. (A) As expected based on previous findings, SNV DNMs are significantly positively correlated with parental age at conception. (B-D) DNMs are not significantly correlated with sex, birth year of the individual for whom we call DNMs, or DNAage of this individual. These comparisons were to confirm that there was no unexpected confounding, and these results are in accordance with predictions.

**Figure S2.**
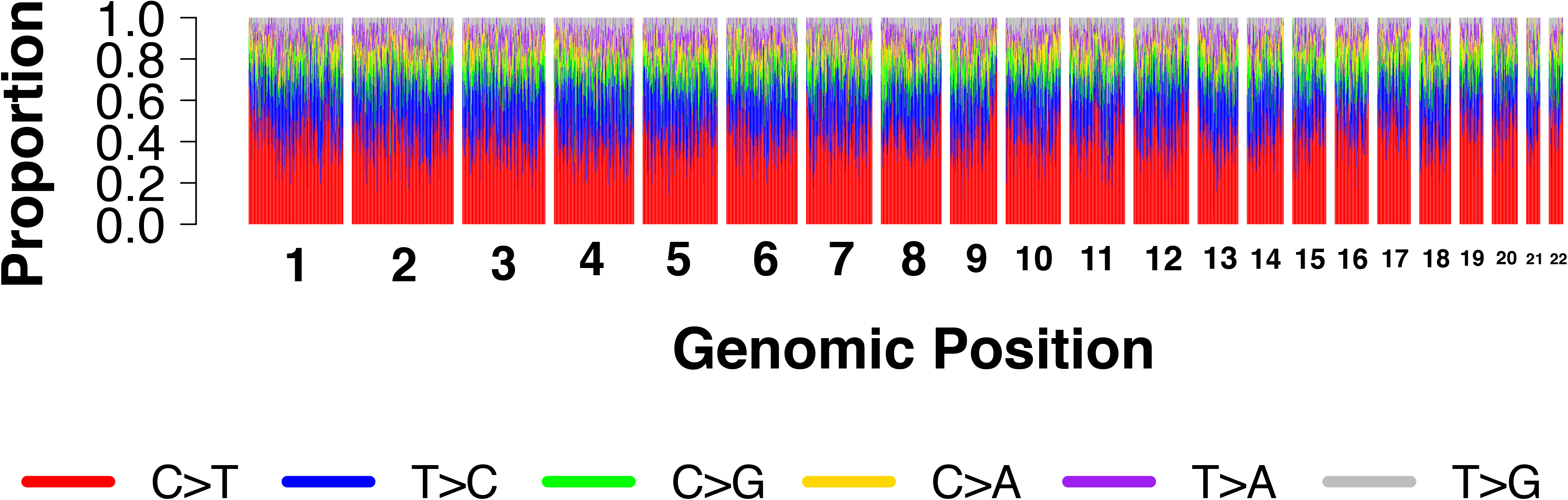
Distribution of 1mer DMNs across the genome. The distribution across the genome for the frequency of each of 6 single base mutation types is shown. These frequencies are estimated for each mutation across each of 2676 1MB windows, and are displayed per window after sorting based on window end position, per autosome.

**Figure S3.**
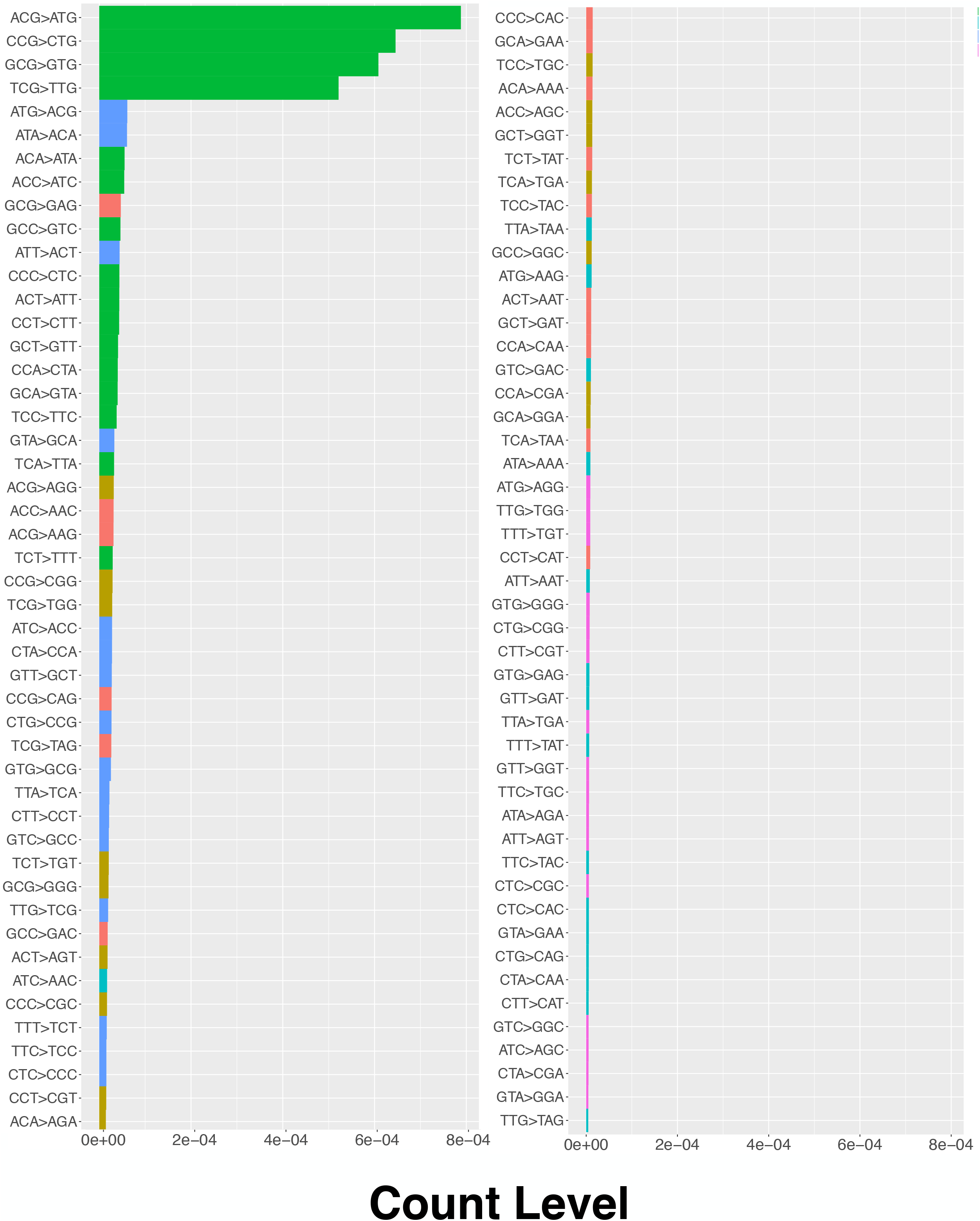
Distribution of 3-mer mutations after normalization by background frequency. Count levels are shown as each 3merDNM count divided by the total count (i.e. frequency) of the 3-mer across the reference genome,which represents a normalization of the count by background frequency. CpG to TpG transitions are much more frequent than the expectation based on 3-mer genome frequency (green, top left). Mutations starting with AT or AC seem to have high frequencies than expected, GCG→GAG are more common than expected based on their center base mutation (i.e. C→A), and TCT→TTT isles frequency than expected based on it center base.

**Figure S4.**
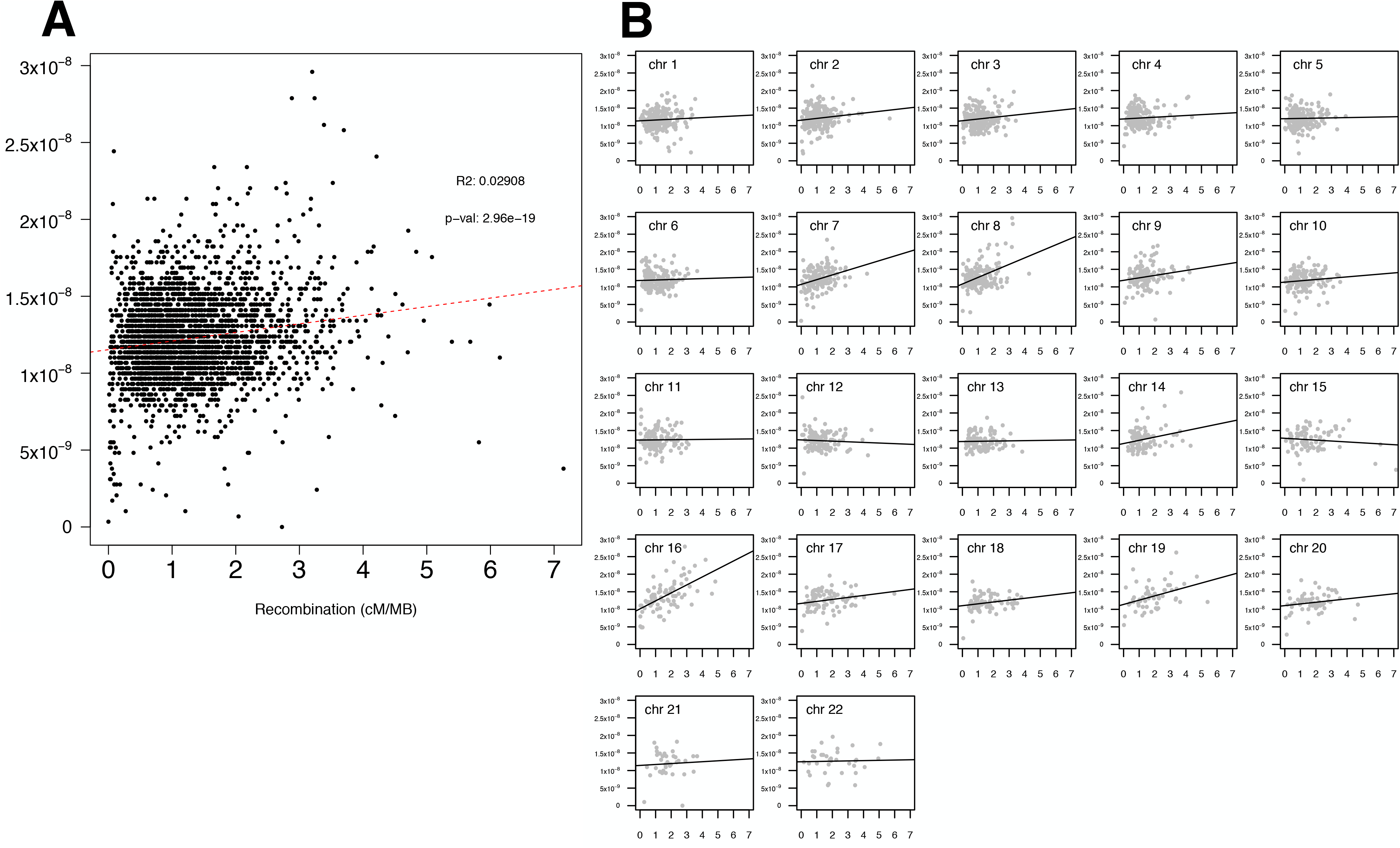
Correlation between SNV DNMs and Recombination Rate. A) Local SNV DNM rates, which were calculated for 2,695 1MB windows across the autosomes, are significantly positively correlated with local recombination rate estimates. B) These correlations vary across the autosomes, with some chromosomes showing strongly significant positive correlations (e.g. chromosomes 7, 8, 9, 16, 19) others showing no correlation (e.g. chromosomes 6, 11, 12, 13, 22), and chromosome 15 showing a significant negative correlation.

**Figure S5.**
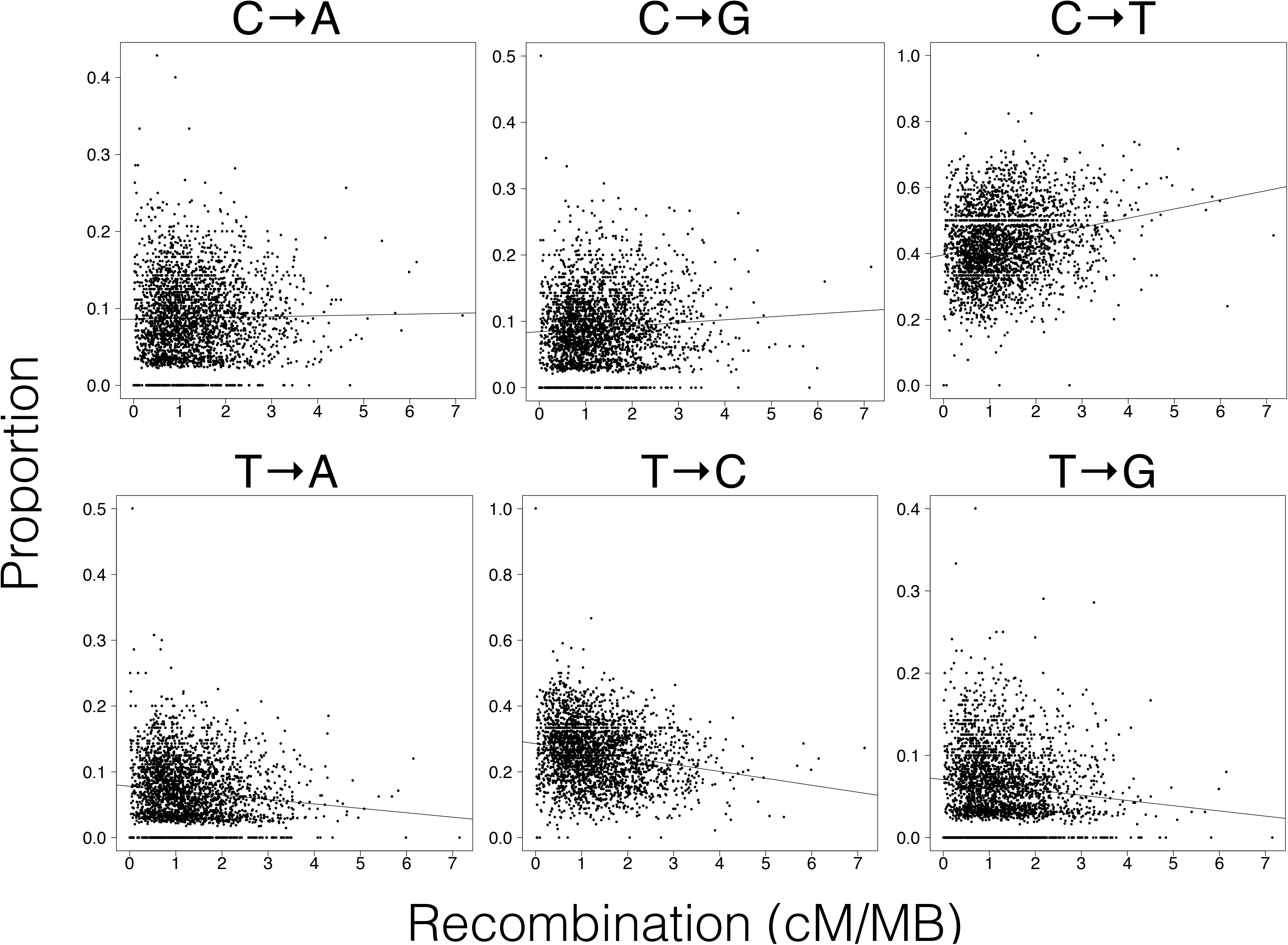
Association of recombination rate and single bas mutation type. C→G and C→T mutation proportions are significantly positively correlated with Recombination rate, while T→A, T→C,and T→G are significantly negatively correlated with recombination. Genomic Location TOPMed Rare Variation Recombination (cM/MB)

**Figure S6.**
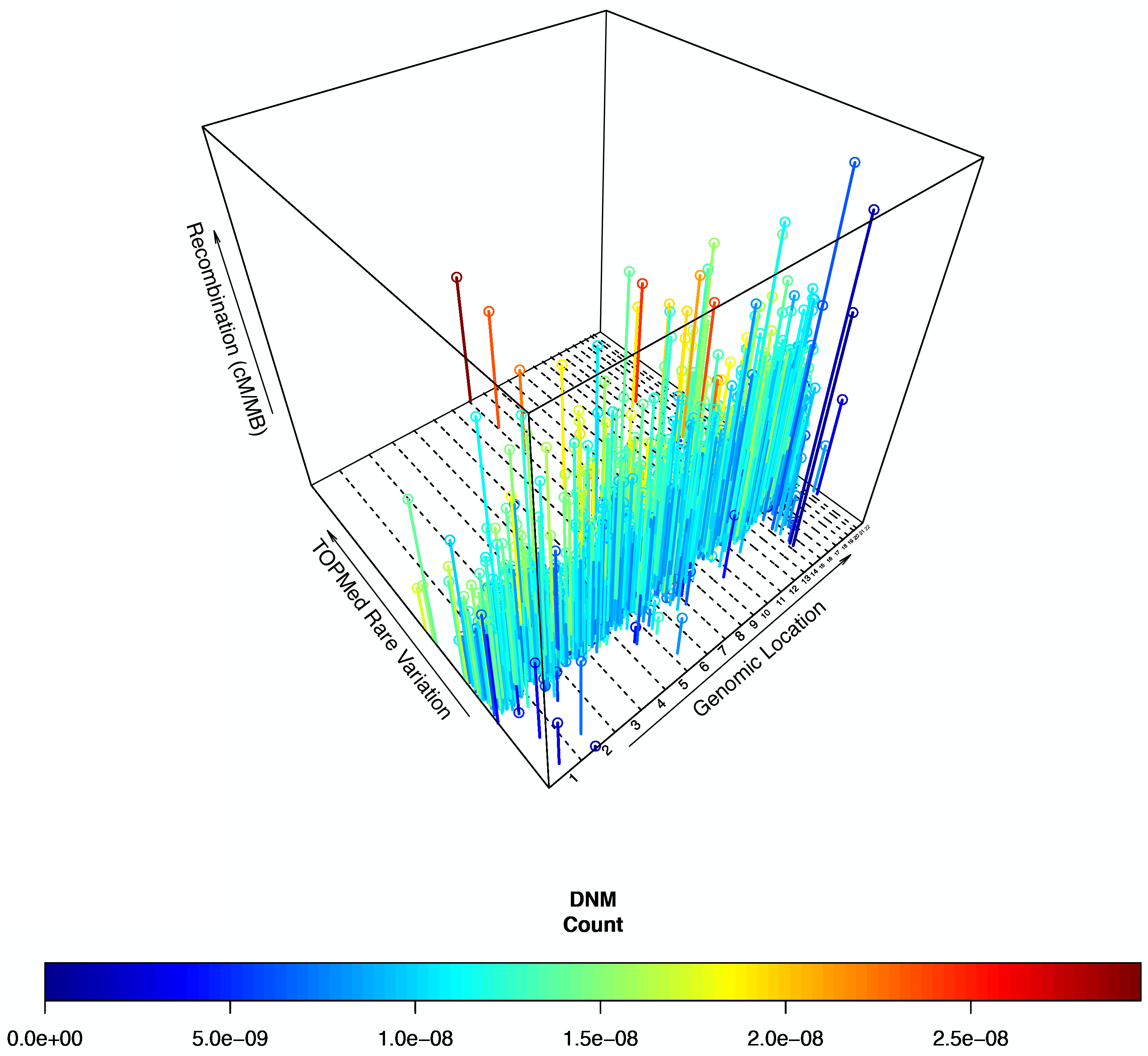
Normalized city plot of rare variation, recombination rate, and DNM rate across the genome. The relationship between DNM rate (blue to red color range), rare variation (Y-axis, ranging from −3.66 to 12.15 Z-scores), and recombination rate (Z-axis, ranging from 0 to 7.77 cM/Mb Y cM/MB) is qualitatively similar across the genome (x-axis, dotted vertical lines divide autosomes 1-22) after adjusting rare variation totals for coding proportion and mappability concerns (see methods). In going from low to high rare variation levels, a blue to red gradient can be seen, as in the unadjusted analysis. This reflects the significant correlation between DNM rate and population-level rare variation. Furthermore, regions with high DNM rates and high variation levels generally have taller bars, which reflects the positive relationship between DNM rate, variation level, and recombination (a few exceptions to this can be seen at the bottom corner of the plot as tall blue bars). Regions with the highest variation levels in the genome, such as those on chromosomes 8 and 16, have the highest DNM rates.

**Figure S7.**
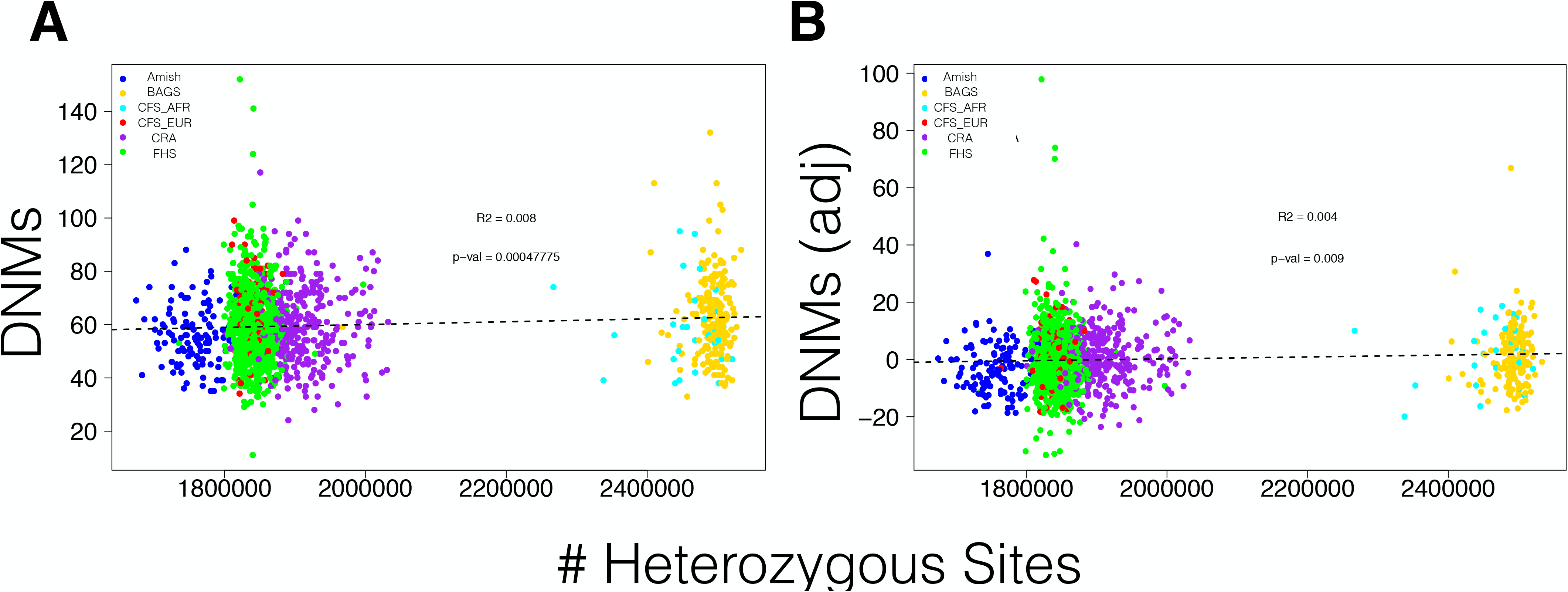
Global Heterozygosity and SNV DNMs. A) Genome Heterozygosity total is significantly but modestly correlated with SNV DNM total, B) and this correlation persists after adjusting for parental age effects.

**Figure S8.**
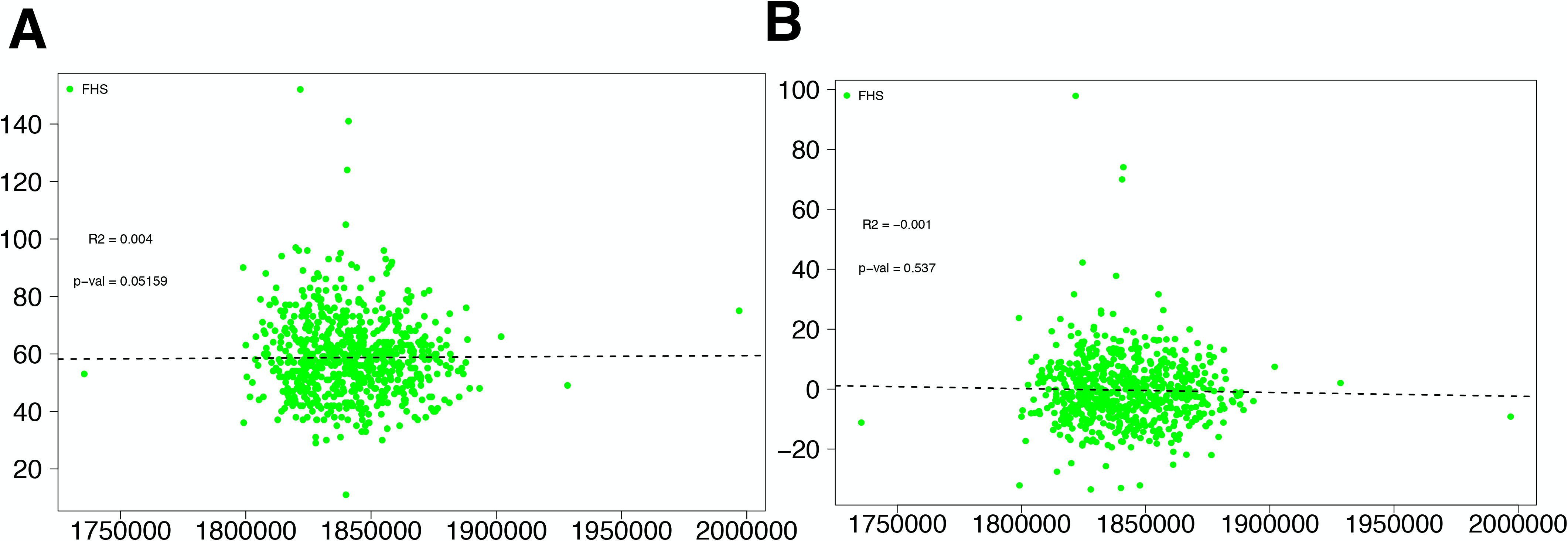
Heterozygosity does not significantly associate with DNMs in FHS. Within the FHS cohort, which represents our largest cohort, there is no correlation between an individual’s total number of heterozygous sites and their number of DNMs.

**Figure S9.**
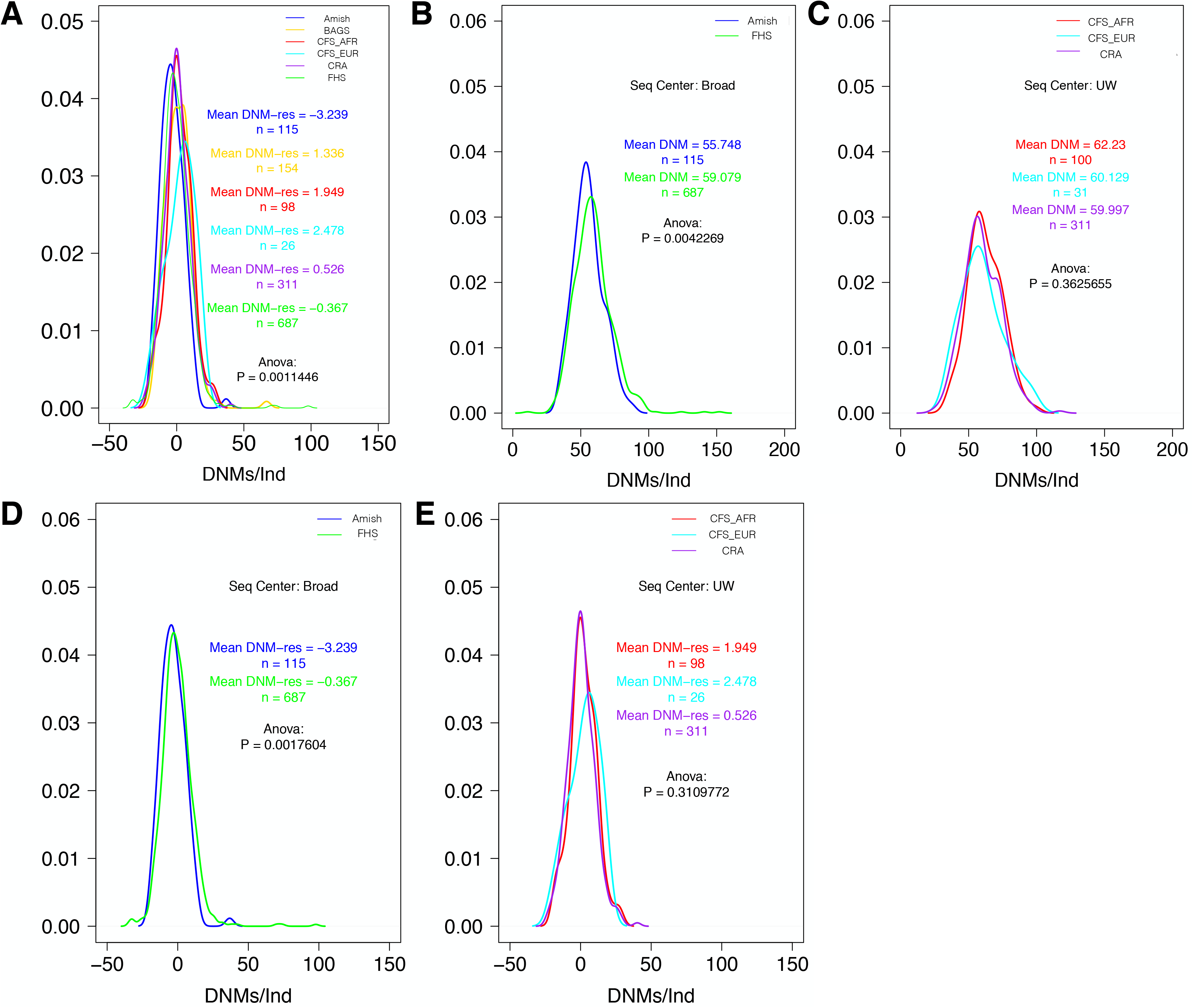
SNV DNM sub-analyses across diverse cohorts. A) After using linear regression to adjust SNV DNM totals for parental age effects, significant differences across cohort persist. B) Comparisons of SNV DNM totals across cohorts sequenced at the same centers show significant differences between individuals Amish (blue) and FHS (green), but C) no significant differences between CFS_EUR, CFS_AFR, and CRA, which represent European, African, and Latino ancestry individuals, respectively. These results persists after adjusting for parental age effects (D and E).

**Figure S10.**
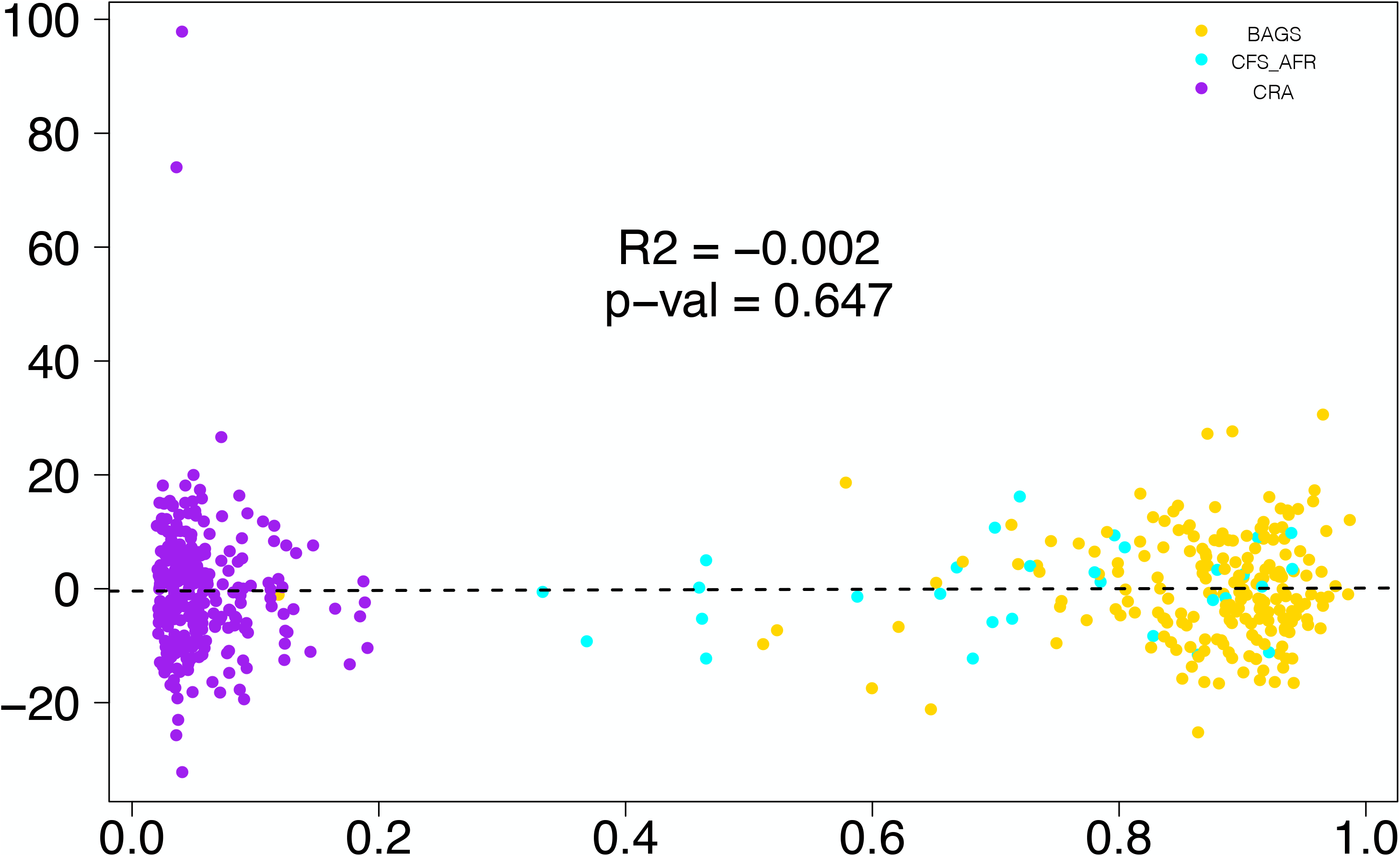
African ancestry proportion does not correlate with SNV DNMs. After using linear regression to adjust SNV DNM totals for parental age effects, there is no correlation between African ancestry proportion (x-axis) and adjusted SNV DNM totals per individual (y-axis).

**Figure S11.**
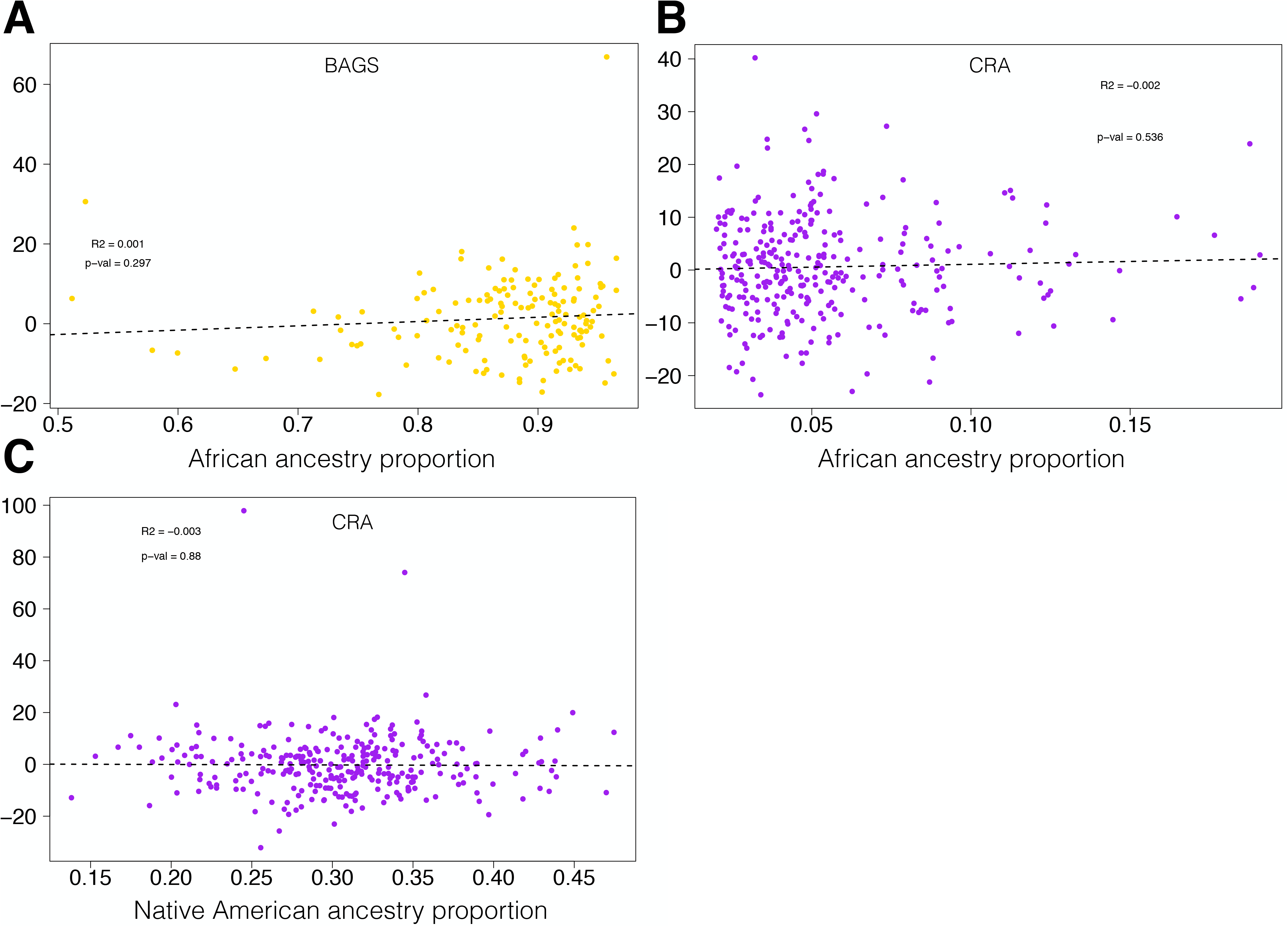
Ancestry proportion does not correlate with SNV DNMs in BAGS or CRA. After adjusting for parental age effects, there are no significant correlations between SNV DNM total per individual and A) African ancestry proportion in BAGS, B) African ancestry proportion in CRA, or C) Native American ancestry proportion in CRA.

**Figure S12.**
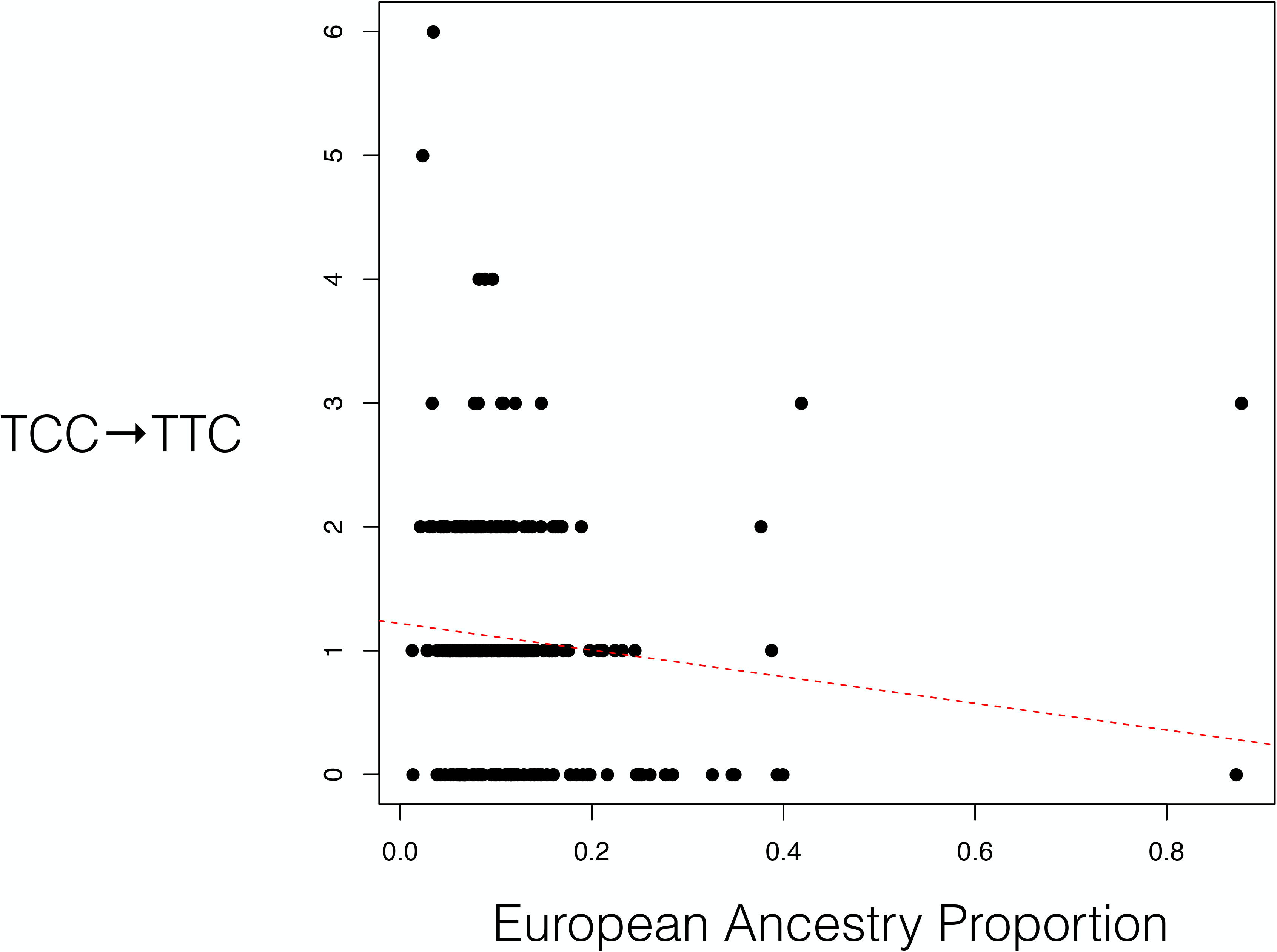
TCC→TTC DNMs and European ancestry in the BAGS cohort. TCC→TTC 3-mer mutation count correlates negatively with global European ancestry proportion within African ancestry individuals from the BAGS cohort (P<6.5×10-3, not significant after Bonferroni correction for multiple testing).

**Figure S13.**
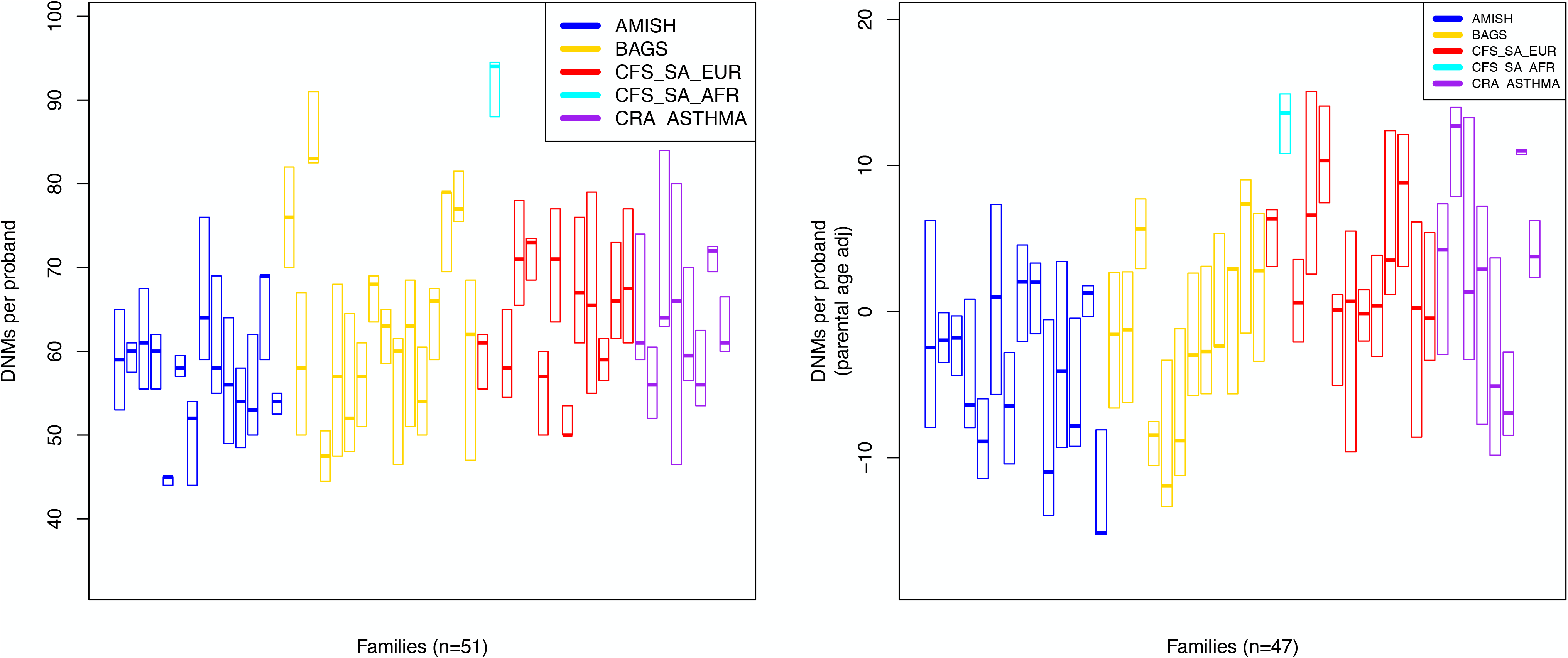
SNV DNMs differ across families. Amongst families with 2 or more children, A) SNV DNM totals per individual are significantly different across family, B) which persist after adjusting for parental age effects (P<0.005).

**Figure S14.**
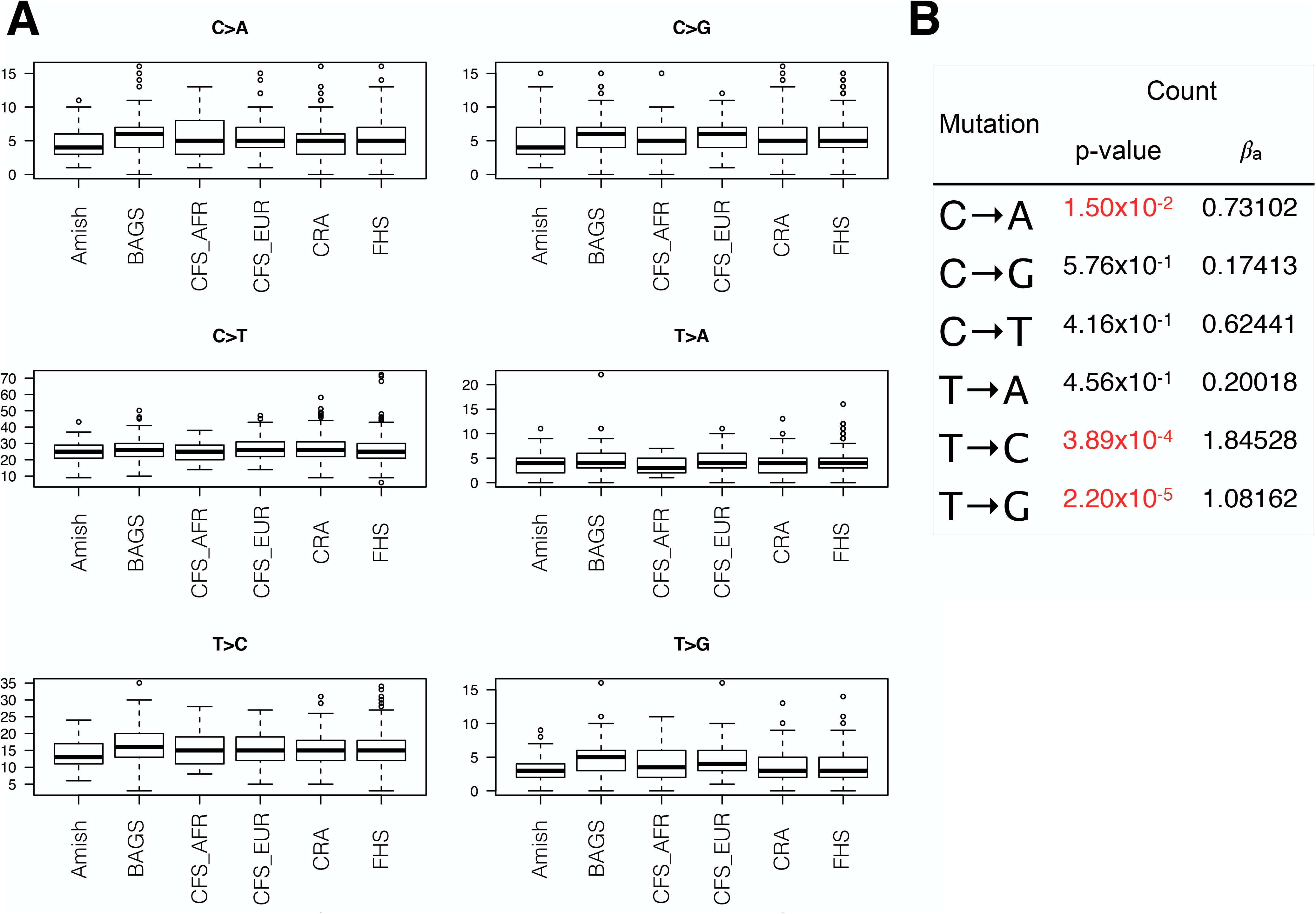
Single base mutation differences across cohort. Significant differences C→A, T→C, and T→G mutations can been seen across cohort (A and B).

**Figure S15.**
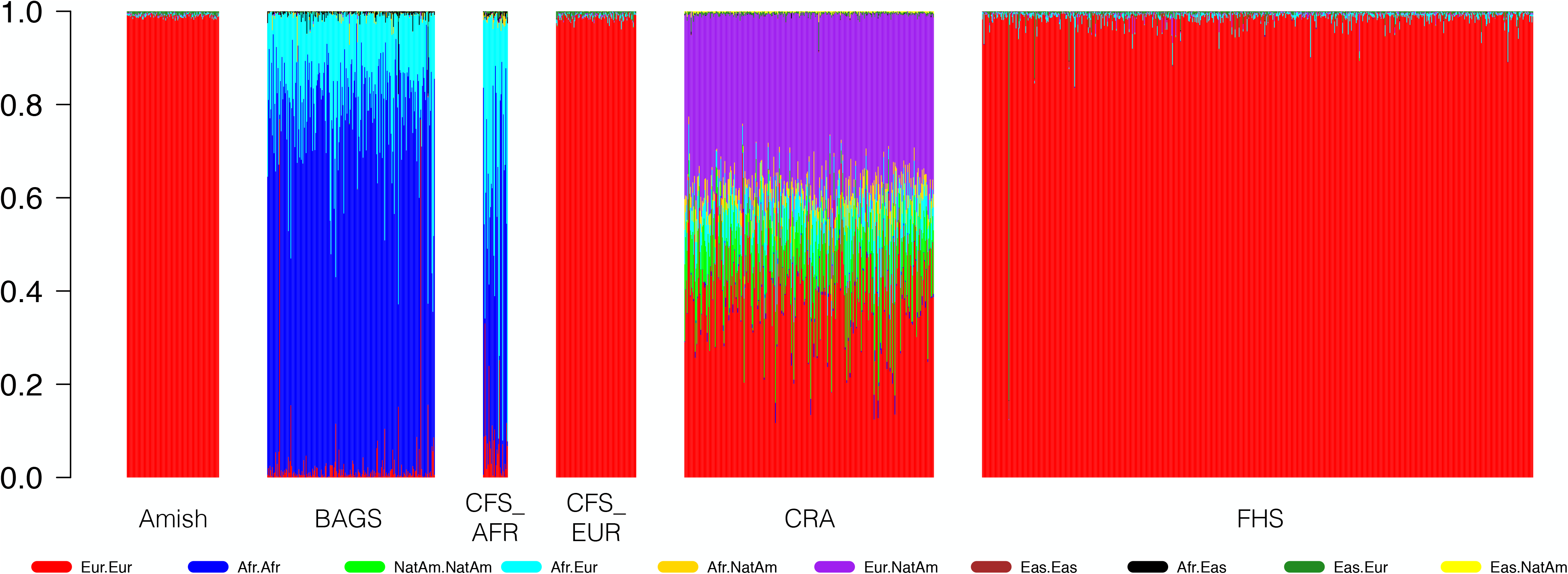
Diplotype admixture proportions across cohorts. The proportion of each diplotype (out of those with appreciable representation in our dataset) is represented by a different color within a stacked bar plot for each sample in our DNM analysis (n=1453). Individuals from the BAGS, CFS_AFR, and CRA cohorts have significant diplotypic admixture, and contain the sampleswe used for our local ancestry analysis.

**Figure S16.**
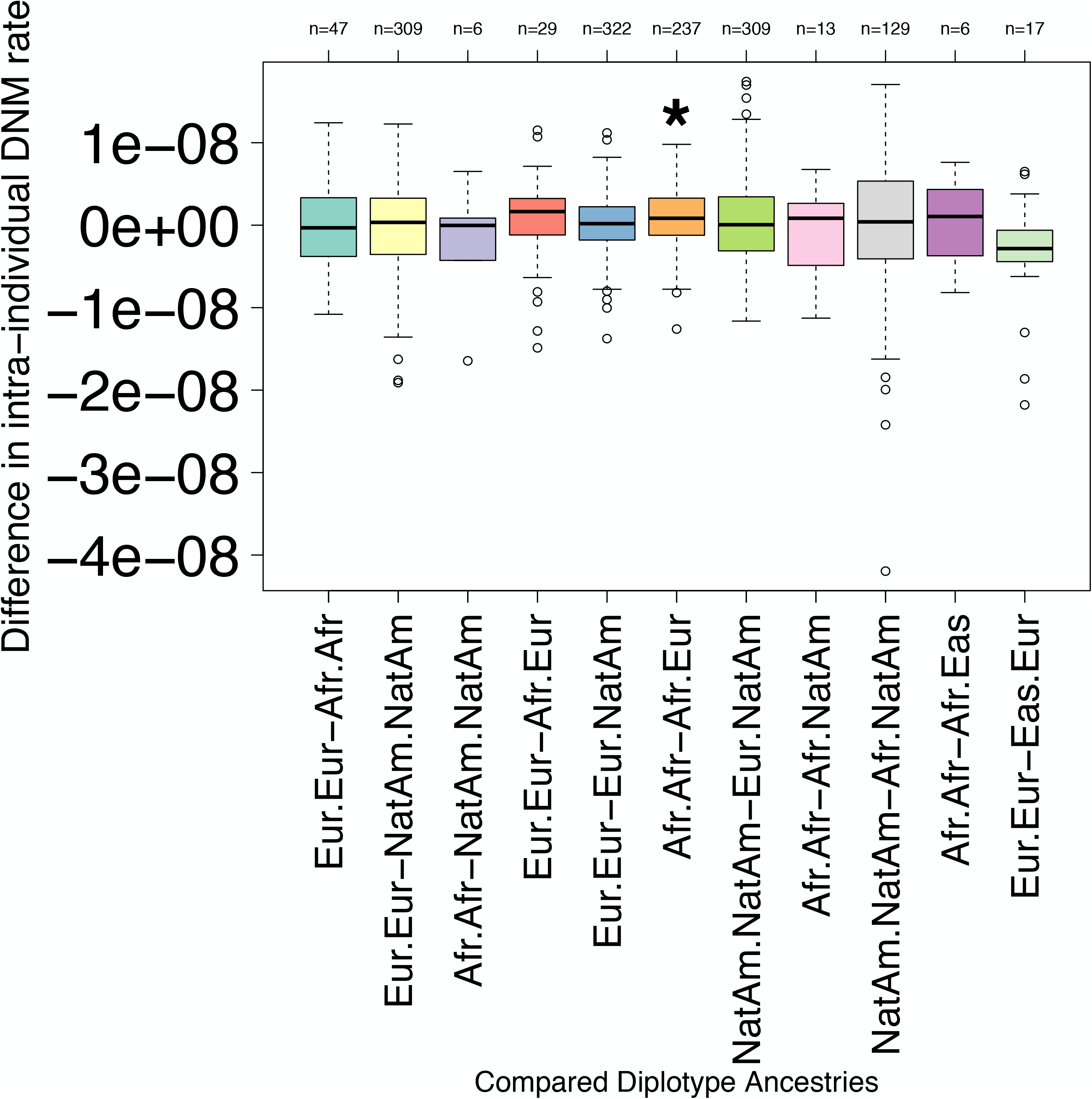
Intra-individual comparisons of mutation rate across ancestry. Every individual who had a minimum of 80 Mb of each of at least two ancestry diplotypes, had individualized DNM rates calculated for each of these diplotypes (i.e. a European/European mutation rate). These individualized rates were then subtracted from one another, and for all pairwise combinations where we had enough representation in our samples (x-axis), we built distributions of these intra-individual DNM rate differences (y-axis). These differences centered around zero, and using paired t-tests, we did not find any consistent significant differences across ancestral background in intra-individual mutation rates. While the Afr.Afr - Afr.Eur (orange) difference just reached significance after multiple testing (P<6.1×10-4, black asterisk), there is no other signal of a higher mutation rate in African ancestry versus European ancestry segments.

